# High Affinity binding of Yeast Nucleolin Nsr1 to Co-transcriptionally Formed G4 DNA Obstructs Replication and Elevates Genome Instability

**DOI:** 10.1101/802876

**Authors:** Shivani Singh, Alexandra Berroyer, Minseon Kim, Nayun Kim

## Abstract

A significant increase in genome instability is associated with the conformational shift of a guanine-run-containing DNA strand into the four-stranded G-quadruplex (G4) DNA. The mechanism underlying the recombination and genome rearrangements following the formation of G4 DNA *in vivo* has been difficult to elucidate but has become better clarified by the identification and functional characterization of several key G4 DNA-binding proteins. Mammalian nucleolin NCL is a highly specific G4 DNA-binding protein with a well-defined role in the transcriptional regulation of genes with associated G4 DNA-forming sequence motifs at their promoters. The consequence of the *in vivo* interaction between G4 DNA and nucleolin in respect to the genome instability has not been previously investigated. We show here that G4 DNA-binding is a conserved function in the yeast nucleolin Nsr1. Furthermore, we demonstrate that the Nsr1-G4 DNA complex formation results in replication obstruction and is a major factor in inducing the genome instability associated with the co-transcriptionally formed G4 DNA in the yeast genome. The G4-associated genome instability and the G4 DNA-binding *in vivo* requires the arginine-glycine-glycine (RGG) repeats located at the C-terminus of the Nsr1 protein. Nsr1 with the deletion of RGG domain supports normal cell growth and is sufficient for its pre-rRNA processing function. However, the truncation of RGG domain of Nsr1 significantly weakens its interaction with G4 DNA *in vitro* and *in vivo* and restores unhindered replication, overall resulting in a sharp reduction in the G4-associated genome instability. Our data suggest that the interaction between Nsr1 with the intact RGG repeats and G4 DNA impairs genome stability by precluding the access of G4-resolving proteins and obstructing replication.

**AUTHOR SUMMARY:** Genome instability is uniquely elevated at sequences containing multiple runs of guanines, which can fold into the unusual, four-stranded G-quadruplex (G4) DNA. In this study, we report a novel finding that a highly conserved G4 DNA binding protein Nsr1 can elevate the rate of recombination and chromosomal rearrangement occurring at a G4 DNA-forming sequence in the genome of *Saccharomyces cerevisiae*. The elevated genome instability requires the C-terminally located RGG domain of Nsr1, which supports the high-affinity interaction between the protein and G4 DNA. The connection between G4-specific genome instability and the function of Nsr1 to form stable complex with G4 DNA led to the hypothesis that the high-affinity Nsr1-G4 DNA complexes can become a barrier to replication. We demonstrate here that the presence of Nsr1 in fact slows the replication past a G4 DNA-containing genomic site and that the RGG domain is required to facilitate such replication block.

## INTRODUCTION

G-quadruplexes or G4 DNAs are unique non-canonical four-stranded DNA structures that can form from guanine-rich nucleic acid sequences [1, 2]. Four guanine molecules interact *via* Hoogsteen bonds to yield G-quartets that in turn stack on top of each other, held together by intervening loops of variable length and sequence to form the G4 DNA. The size and orientation of the loops can determine the relative stability of various G4 DNA configurations [3]. Though some computational analyses revealed that >375,000 and >1,400 G4 motifs are in the human and the *Saccharomyces cerevisiae* genomes, respectively, it is still unclear how many of these QFP sequences actually assume the stable G4 configuration *in vivo* [4, 5]. Nevertheless, it is becoming quite evident that these structural transformations potentially provide a substantial informational capacity to DNA with specific functions [1]. Recent bioinformatic studies in human, yeast and bacterial genomes have revealed that G4 motifs are not randomly located within genomes, but particularly enriched in certain functional regions, such as those in ribosomal and telomeric DNA, G-rich micro- and minisatellites and mitotic and meiotic double-strand break (DSB) sites [6–8]. Their high prevalence in and around genes particularly proximal to transcriptional start sites (TSSs) suggests a possible role as *cis*-acting regulatory elements [9–11].

The presence of G4 DNA was recently shown to impair DNA replication by hindering the progress of replicative polymerases, and the failure to resolve these structures transforms the sequence motifs into potential hotspots for genomic instability [12]. In human, the occurrence of G4 DNA motifs reportedly overlap with recombination-prone regions such as certain proto-oncogenes and the sites of frequent translocation breakpoints [13, 14]. Consistently, chromosomal translocations in the proximity of G4 motifs have been observed in a subset of leukemias and lymphomas [15]. The potential G4 DNAs additionally have been correlated with a number of human neurological diseases, such as the frontotemporal dementia (FTD), amyotrophic lateral sclerosis, Alzheimer’s and fragile X syndrome [16–18]. The resolution of G4 DNA structures is thus imperative in preserving genome integrity.

The genetic tractability of *S. cerevisiae* has provided considerable insight into the mechanisms involved in maintaining stability of G-rich repetitive sequences. The instability at the G-rich human minisatellite CEB1 inserted in the *S. cerevisiae* genome was shown to be dependent on the ability of the CEB1 motif to form G4 DNA and was not observed with other tandem repeats lacking G4 DNA-forming potential [19, 20]. The G4-forming sequence derived from the guanine-run containing immunoglobulin switch Mu (Sμ) region becomes highly unstable when actively transcribed in the context of the yeast genome. Transcription conferred a critical strand bias since genome rearrangements at Sμ were elevated only when the guanine-runs were located on the non-transcribed strand [21]. The direction of replication and transcription, when in a head-on orientation, further elevated genome instability at the Sμ sequence [22]. Also, at the Sμ sequence, the lack of functional topoisomerase 1 (Top1) significantly elevated various types of genome instability, likely by facilitating the structural transition of a G-rich sequence to a G4 structure due to the accumulation of negative helical stress in DNA. The loss of heterozygosity and copy number alterations (deletions and duplications), both of which are frequently observed in cancer genomes, were also elevated when the Sμ sequence was actively transcribed.

The biological functions of G4 DNA are largely dependent on the protein factors that modulate the G4-conformation and/or serve as a bridge to recruit additional protein regulators [23]. These G4 binding proteins can be classified into three functional groups: 1) telomere-related proteins, such as the shelterin complex, human CST (CTC1-STN1-TEN1), and yeast Rap1 and Est1 [24–26]; 2) proteins that unfold and/or process the G4 structure, such as the helicases including RecQ family helicases hBLM, hWRN, ySgs1, and yPif1 [27]; and 3) proteins that stabilize G4 structures including MAZ and nucleophosmin [28, 29]. Mutations in some of these G4-interacting proteins have been linked to genetic diseases such as Werner syndrome, Fanconi anemia, and cancer [27, 30–32]. A defective BLM helicase failed to unwind G4-DNA and caused increasing recombination frequencies and a high incidence of cancer in Bloom’s syndrome [33]. More recently, the co-transcriptional activator Sub1, which interacts with both G4 DNA and the G4-helicase Pif1, was shown to suppress the G4-associated genome instability by facilitating the recruitment of Pif1 helicase to co-transcriptionally formed G4 DNA structures [34].

The human Nucleolin (NCL) is a highly abundant and conserved nucleolar phosphoprotein. Its major function is in the ribosomal RNA maturation with additional roles in chromatin remodeling, transcription, and apoptosis [35]. The altered expression and subcellular localization of NCL is a common biomarker of a variety of cancers demonstrating its clinical relevance [3, 36–38]. Although initially described as a G4-RNA binding protein, recent evidences indicate that NCL preferentially and selectively binds both endogenous and exogenous G-rich sequences that can fold into G4 DNA. It is suggested that NCL acts as a chaperone to promote the correct folding of complex nucleic acid structures [39–41]. Together with the nuclear riboprotein hnRNPD, NCL forms a lymphocyte-specific complex LR1 (lipopolysaccharide responsive factor 1), which binds at the G4 DNA-forming Immunoglobulin heavy chain (IgH) switch regions [42]. The binding of NCL to the G4-forming hexanucleotide repeat expansion (HRE) (GGGGCC)_*n*_ in *C9orf72* has been reported to be responsible for the initiation of molecular cascades that lead to neurodegenerative diseases [17]. NCL is composed of three main structural domains; the amino-terminal domain containing four acidic stretches was shown to induce chromatin decondensation through interaction with histone H1 [43], while the central region containing tandem RNA binding domains (RBDs) and the multiple RGG (arginine/glycine/glycine) boxes at the C-terminal domain contribute to its high-affinity interaction with G4 DNA [44, 45].

Similar to the human protein, the yeast Nucleolin Nsr1 has been demonstrated to be involved in pre-rRNA processing and ribosome production [46, 47]. While there are no significant sequence similarities in the amino-terminal regions between yNsr1 and hNCL, the C-terminal half of yNsr1, consisting of two tandemly repeated putative RNA binding domains (RBDs) and the multiple RGG (arginine/glycine/glycine) motifs, has a high sequence similarity to the carboxyl-terminal part of hNCL (37% identity in 249 amino acids) [46, 48]. Unlike the plethora of studies highlighting the NCL/G4 interaction, the role of yeast Nsr1 related to G4 DNA has not been extensively studied. In the current study, we examined whether the yeast Nsr1 has a role in the structural flux of G4-forming sequences. We identified an important biological role of Nsr1 in enhancing various types of genome rearrangements associated with the co-transcriptionally formed G4 DNA. We show that the yeast Nsr1 can bind G4 DNA both *in vitro* and *in vivo* and that the disruption of Nsr1 lowers the G4-associated recombination, while its overexpression further exacerbates instability in a dose-dependent manner in cells lacking Top1. The C-terminal RGG domain of Nsr1 is required to form a complex with G4 DNA *in vivo*, to obstruct replication, and to promote genetic instability, but is dispensable for the rRNA processing function of Nsr1. Our results point to an important role of Nsr1 in G4-associated genome maintenance.

## RESULTS

### Nsr1 enhances G4-mediated recombination in cells lacking Topoisomerase 1

To understand the elevated genome instability associated with G4-forming sequences, we previously developed a recombination reporter assay in the yeast model system [21]. In this reporter assay, a model G4-forming sequence from the murine immunoglobulin heavy chain switch Mu region (Sμ) is inserted into the yeast genome within the *LYS2* gene under the control of a tetracycline/doxycycline-repressible promoter (*pTET*). The Sμ sequence was inserted either in the physiological (GTOP) or in the inverted orientation (GBTM), placing the G-runs in the non-transcribed strand (NTS) or in the transcribed strand (TS) respectively. The formation of G4 DNA is favored when the G-rich strand located on the NTS is transformed into a single stranded state during transcription, freeing the guanine bases to interact with each other through Hoogsteen base-pairing. When G-runs are located in the TS, they will be occupied in base pairing with the nascent RNA strand and will not be free to fold into G4 DNA. Thus, any factors involved in the formation or stability of G4 DNA should affect the recombination occurring at the *pTET-lys2-GTOP* construct, with little to no effect on the rate of recombination at the *pTET-lys2-GBTM* construct [21, 22, 34].

To determine whether Nsr1 plays a role in the G4-mediated genome instability, we deleted the *NSR1* gene in strains containing the *pTET-lys2-GTOP or -GBTM* construct and checked the rate of recombination at this locus. In the *WT* background, the deletion of *NSR1* resulted in a slow growth phenotype of the cells as previously reported [47]. Under high transcription conditions, the rates of recombination at the *pTET-lys2-GTOP* or *-GBTM* cassette in *nsr1*Δ strains were not changed from those in *WT* strains (Figure 1A).

**Figure 1.**
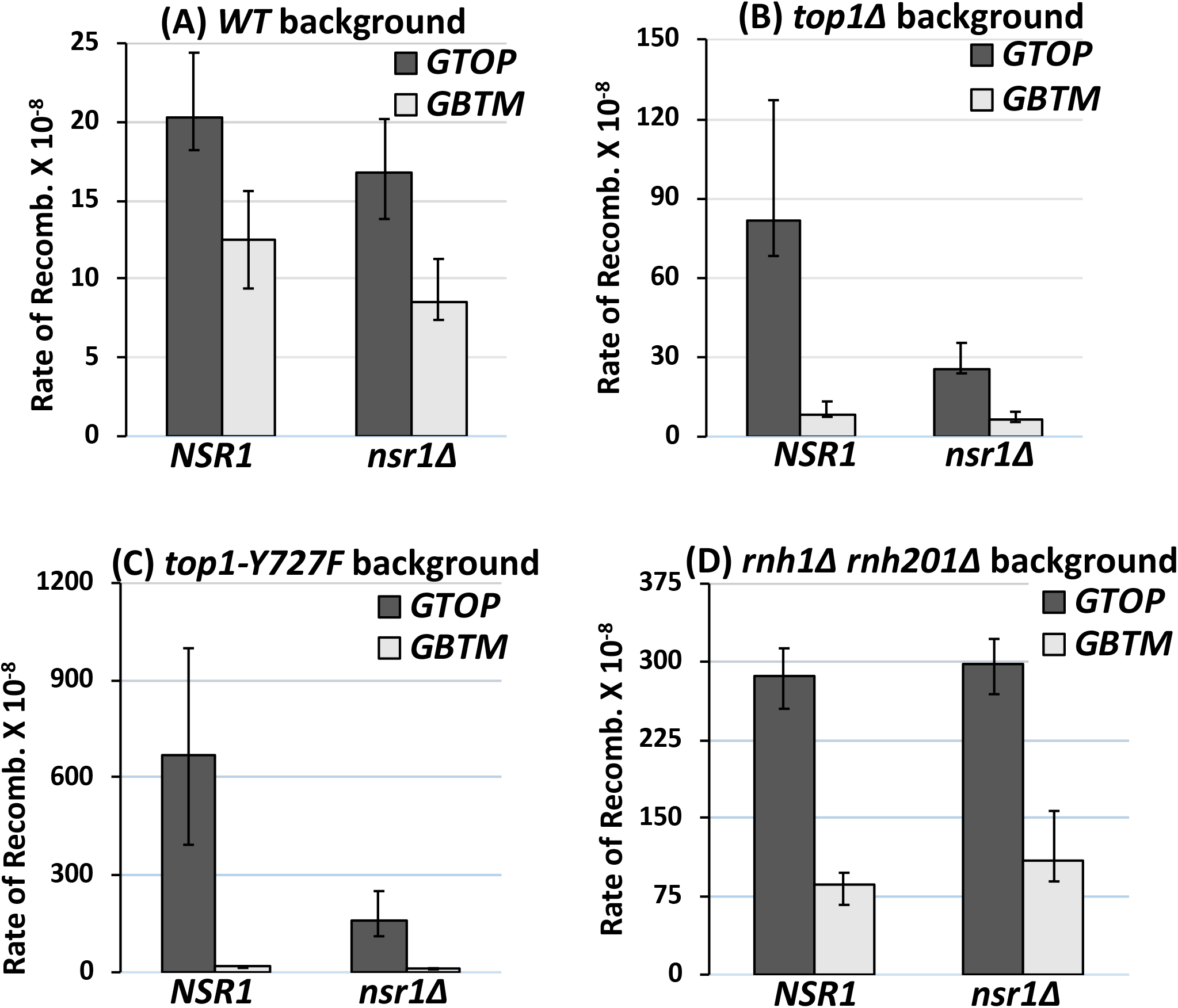
The rates of G4-associated recombination in *nsr1*Δ strains. Guanine-runs are on the non-transcribed, top strand in a single stranded state in *pTET-lys2-GTOP* cassette and on the transcribed strand annealed to the nascent RNA in the *pTET-lys2*– *GBTM* cassette. All graphs are showing the rates of recombination (×10^−8^). Error bars indicate 95% confidence intervals. (**A**) Recombination rates in *WT* backgrounds. (**B**) Recombination rates in *top1*Δ backgrounds. (**C**) Recombination rates in *top1Y727F* backgrounds. (**D**) Recombination rates in *rnh1Δ rnh201*Δ backgrounds.

Topoisomerase I (Top1) was previously identified to be a crucial factor for G4 mediated genome instability in yeast [21, 22, 49]. Accumulation of negative supercoils in *top1*Δ yeast cells was shown to promote recombination at the *pTET-lys2-GTOP* construct. In order to determine whether Nsr1 modulates the G4-associated genomic instability in the absence of Top1, we deleted the *NSR1* gene in a *top1*Δ background. The rate of recombination for the *pTET-lys2-GTOP* was reduced by ∼5-fold in the *top1*Δ *nsr1*Δ strain compared to *top1*Δ (Figure 1B). For the *pTET-lys2-GBTM*, the rates of recombination were indistinguishable between *top1*Δ and *top1*Δ *nsr1*Δ backgrounds. When the transcription from the *pTET* promoter was repressed by the addition of 2μg/ml doxycycline in the media, the deletion of *NSR1* did not affect the rates of recombination in *WT* and reduced slightly in *top1*Δ background (Figure S1).

The catalytically inactive mutant of Top1 (Top1-Y727F) with the mutation of catalytic tyrosine to phenylalanine results in complete ablation of its function in removal of supercoils, while not affecting its DNA binding activity [50]. Possibly due to its high G4-binding ability, the expression of Top1-Y727F results in recombination rates that are ~4-fold higher than in the absence of Top1 (*top1*Δ) [49]. When Nsr1 was disrupted in the *top1Y727F* backgrounds, the recombination rate at the *pTET-lys2-GTOP* cassette was significantly decreased and was comparable to the rate in the *top1*Δ strain (Figure 1C). The disruption of Nsr1 did not affect the rate of recombination at the *pTET-lys2-GBTM* cassette.

### Nsr1 does not affect the G4-associated recombination in RNase H-deficient cells

Disrupting both RNase H1 and RNase H2 in yeast leads to the accumulation of transcription-associated RNA:DNA hybrids or R-loops [51] and subsequently elevates levels of recombination for both the *pTET-lys2-GTOP and -GBTM* constructs [21]. The G-loop, a higher order structure consisting of R-loop and G4 DNA, could arise from either the enhanced negative supercoils leading to G4 formation upon disruption of Top1 or due to the failure to degrade RNA leading to R-loops upon disruption of RNase Hs. Therefore, we checked whether, in addition to reducing recombination rates in a *top1*Δ strain, the deletion of Nsr1 could affect the R-loop mediated enhanced recombination rates. In the triple deletion mutants *rnh1*Δ *rnh201*Δ *nsr1*Δ, there was no significant alteration in the rate of recombination for the *pTET-lys2-GTOP or - GBTM* construct (Figure 1D). This data suggests that Nsr1 induced genomic instability is specific to G4 DNA formed due to the enhanced negative helical torsion under high transcription and not due to the RNA: DNA hybrid accumulation.

### Nsr1 promotes gross chromosomal rearrangements in the *top1*Δ backgrounds

We previously reported that, in the absence of functional Top1, the co-transcriptionally formed G4 DNA at the *pTET-lys2-GTOP* construct leads to the increase in ectopic recombination as well as in the gross chromosomal rearrangements (GCRs) [22]. In order to measure the rates of GCR, we used a modified form of the GCR reporter system developed by Chen and Kolodner [52]. In this reporter system, the *URA3* gene was integrated into the left arm of chromosome V (CHR5) replacing the *HXT13* gene located ∼8.5 kb centromere-distal to the *CAN1* gene. The *pTET-lys2-GTOP* or -*GBTM* cassette containing the Sμ G4 motif, as described before [22], is integrated immediately centromere-proximal to *CAN1.* The loss of functional *CAN1* or *URA3* results in resistance to the drug canavanine (Can) or 5-Fluoroorotic acid (5-FOA), respectively. Thus, by simultaneous selection against two counter-selectable markers (*URA3* and *CAN1*), complex genome rearrangement can be measured. Using this modified GCR assay, it was previously shown that in the high transcription conditions, the disruption of Top1 leads to a significantly higher (∼30-fold) GCR rate for *pTET-lys2–GTOP*, where guanine-runs are present on the NTS, compared to the *pTET-lys2–GBTM* construct where guanine-runs are on the TS [22]. When we deleted NSR1 in *WT* backgrounds, there was no change in the GCR rates for *pTET-lys2–GTOP* or *-GBTM* (Figure 2A). However, the disruption of Nsr1 in *top1*Δ backgrounds resulted in a significant, ~6-fold reduction of the GCR rates for the *pTET-lys2-GTOP* but not for *-GBTM* construct (Figure 2B). We further assessed whether the function of Nsr1 is specific to the Sμ G4-mediated genome instability or extends to other G4 motifs as well. In the GCR reporters modified to contain the G4 motif from TCF3 translocation breakpoint [53], *NSR1* deletion by itself did not alter the GCR rates. But, in a similar manner to the GCR reporter containing Sμ G4, when Nsr1 was disrupted in a *top1*Δ strain, the GCR rates were severely lowered by ~9-fold at the *pTET-lys2-GTOP (TCF3)* construct, while at the pTET*-lys2-GBTM (TCF3)* construct the GCR rates were not significantly changed (Figure 2C and 2D). Overall, these data indicate that Nsr1 promotes the instability associated with the co-transcriptionally formed G4 DNA.

**Figure 2.**
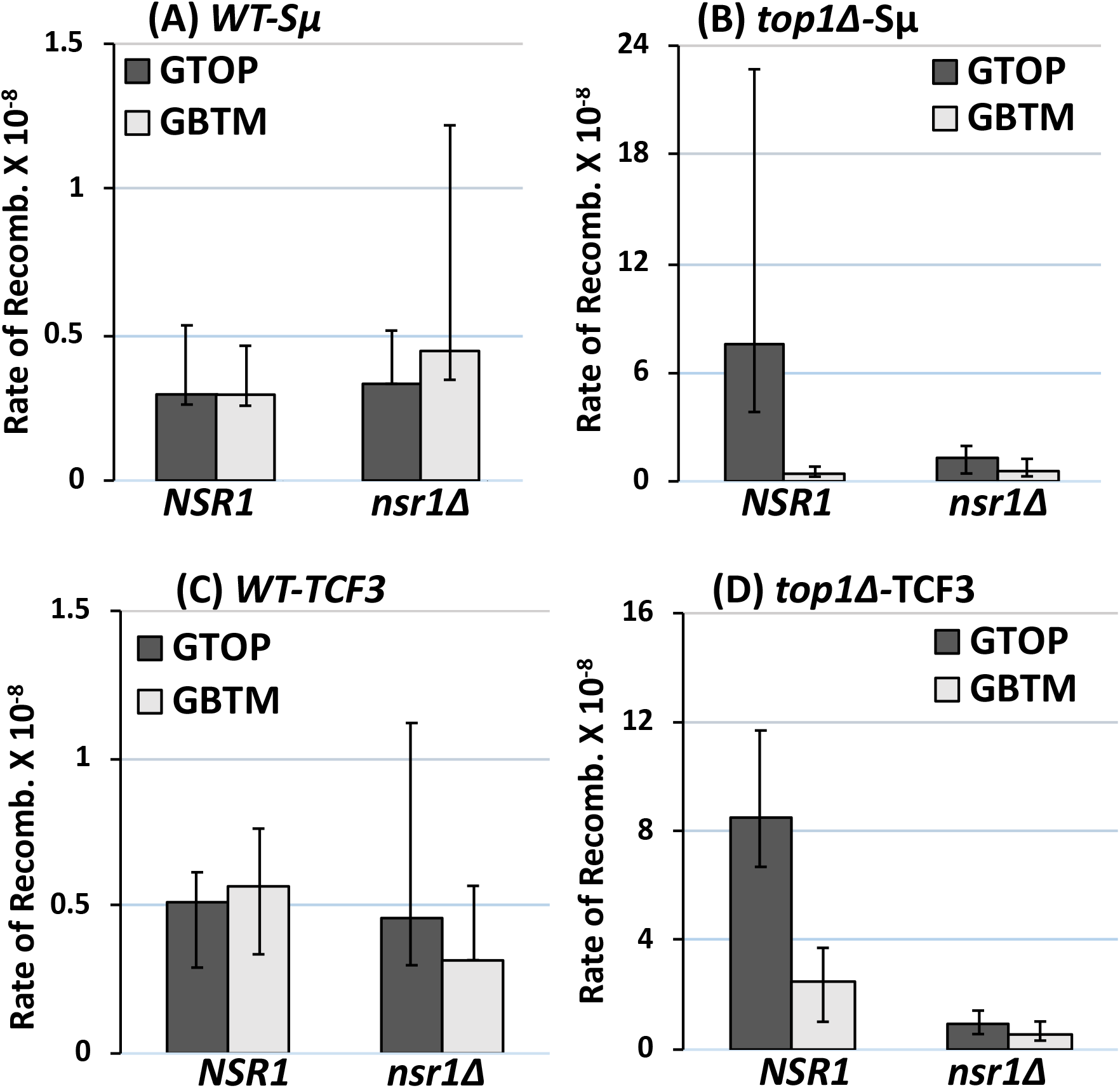
Rates of Gross Chromosomal Rearrangement (GCR) in *nsr1*Δ strains. All graphs are showing the rates of 5-FOA^r^/CAN^r^ or GCRs (×10^−8^). The rates of GCRs occurring at CHR5 containing the *pTET-lys2-GTOP* or *–GBTM* cassette are determined by the method of median. 95% confidence intervals are indicated by the error bars. GCR rates in strains with the reporter cassette containing the Sμ-G4 in (**A**) *WT* backgrounds (**B**) *top1*Δ backgrounds. GCR rates in strains with the reporter cassette containing TCF3-G4 in (**C**) *WT* backgrounds (**D**) *top1*Δ backgrounds.

### The RGG domain of Nsr1 is necessary for the elevated G4-associated recombination but not for the support of normal cell growth or pre-rRNA processing

For the complementation study, a full length *NSR1* gene construct along with its own promoter was cloned into the centromeric plasmid pRS316. Since the C-terminal RGG domain of Nucleolin has been shown to be important for inducing and stably binding G4 structures [40, 44], we also cloned the N-terminal 1-350 residues of Nsr1 lacking the RGG domain (Nsr1ΔRGG) (Figure 3A). The expression of both full length Nsr1 and Nsr1ΔRGG was confirmed using anti-Nsr1 antibody (Figure S2A). The ectopic expression of full length Nsr1 in the *top1*Δ *nsr1*Δ double mutant strain elevated the rate of recombination for the *pTET-lys2-GTOP* by 8.5-fold, thereby resulting in the rates of recombination that were similar to a *top1*Δ transformed with the empty vector (Figure 3B and see Figure 4B). However, the expression of Nsr1ΔRGG did not elevate the rate of recombination at the *pTET-lys2-GTOP* cassette (Figure 3B). The rates of recombination of the *pTET-lys2-GBTM* cassette were not affected by either Nsr1- or Nsr1ΔRGG-expression. We next expressed the N-terminal PASK region of Nsr1 (1-171a.a.; N-term Nsr1 in Figure 3A), or the C terminal region (C-Term; 172 – 414a.a.). The N-term construct was unable to complement for the loss of Nsr1 in enhancing G4-induced recombination (Figure 3C). However, the C-terminal deletion construct elevated the rates of recombination albeit not as robustly as the full length Nsr1. The rates of recombination for the C-terminal deletion construct were 1.8-fold higher than the rates of recombination for the empty vector. These results suggest that the RBD and RGG domains at the C-term of Nsr1 are required for promoting G4 associated instability.

**Figure 3.**
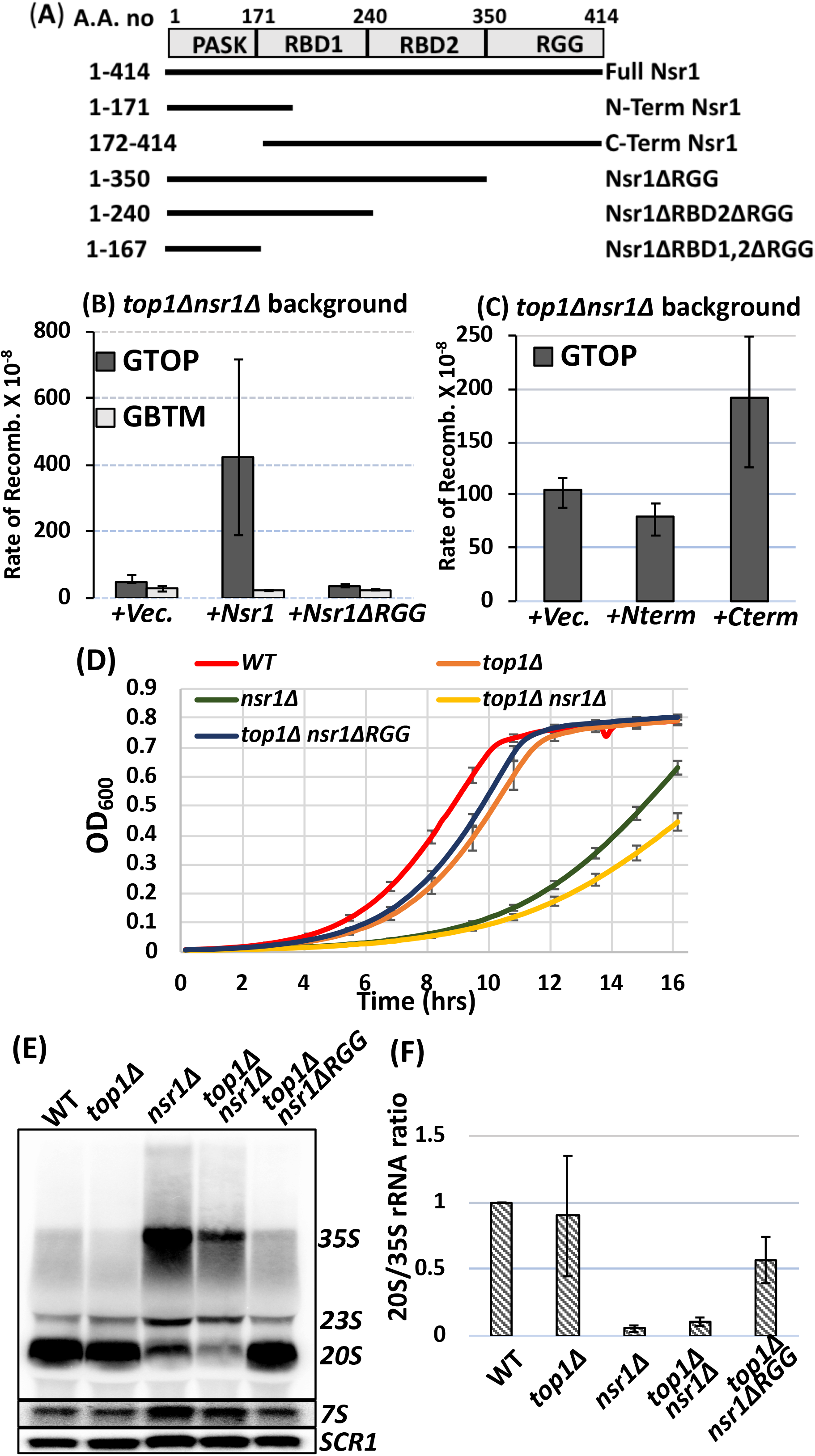
Complementation of NSR1-deletion. (**A**) Domain organization of Nsr1. The deletion constructs used for complementation are indicated. **(B)** and **(C)** Graphs are showing the rates of recombination (×10^−8^). Error bars indicate 95% confidence intervals. The *top1*Δ *nsr1*Δ double mutant was transformed with empty vector (*+Vec*), Nsr1-expression plasmid (*+Nsr1*) or the indicated deletion plasmids followed by fluctuation analysis to determine the rates of recombination. (**D**) Effect of *nsr1*Δ deletion on growth rate. Overnight grown cells from indicated strains were grown to log phase in liquid YEPD media at 30 °C. Cells were then diluted ~1X10^6^ cells/ml in fresh medium. Cell growth at 30 °C was monitored by measuring OD_600_. The data represent averages of eight cultures with the error bars indicating the standard deviation. (**E**) Total RNAs were isolated from the depicted strains followed by northern blot probing with labeled oligos. The *SCR1* RNA serves as a loading control. (**F**) Ratio of 20S/35S in indicated strains normalized to the *WT* ratio.

**Figure 4.**
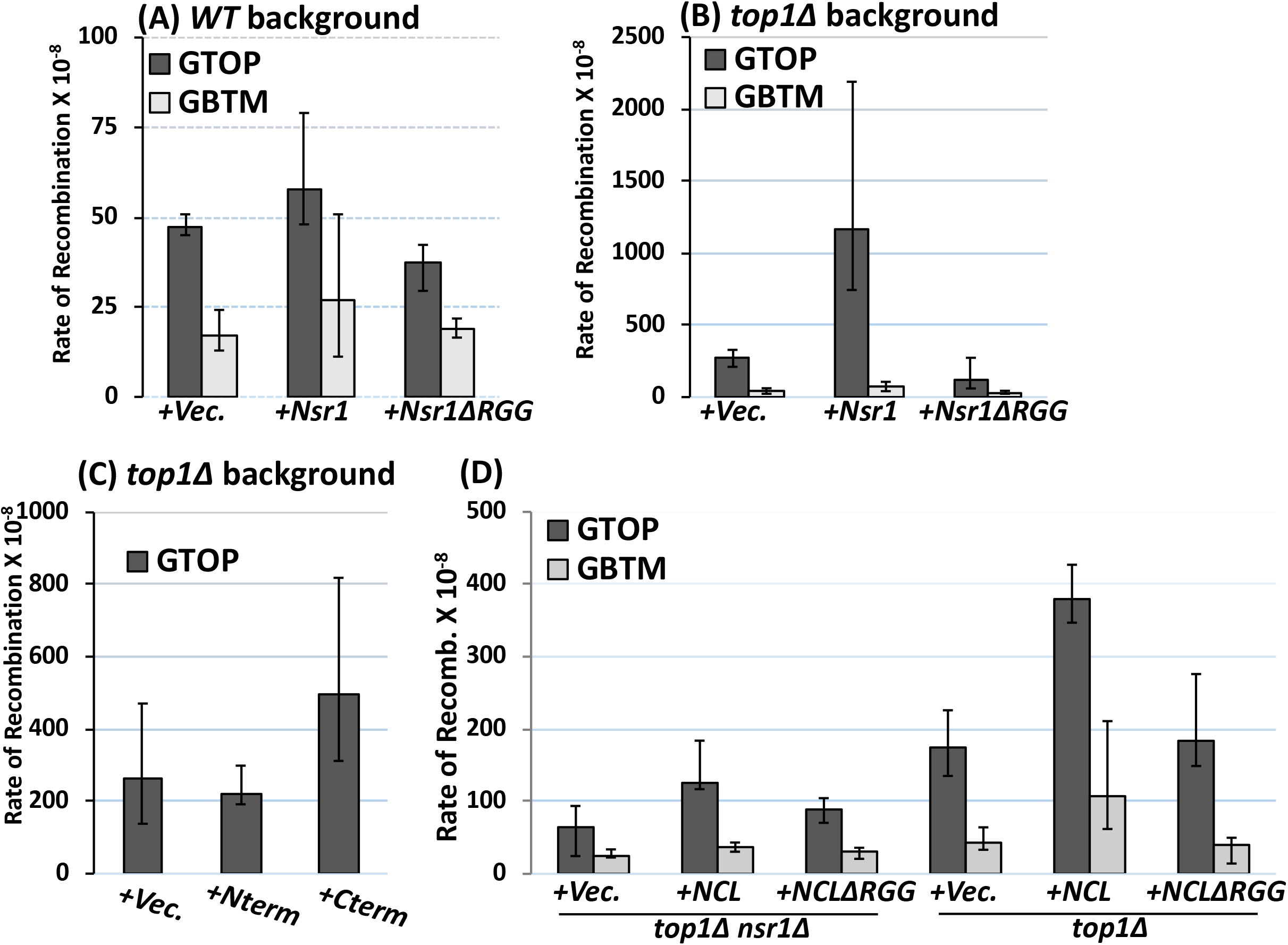
Overexpression of Nsr1 or hNCL. Overexpression of the full length or various deletion constructs of Nsr1 in **(A)** *WT* background, and **(B** and **C)** *top1*Δ background. **(D)** The *top1*Δ *nsr1*Δ or *top1*Δ yeast cells were transformed with empty vector (*+*Vec), hNCL-expression plasmid (*+*NCL) and hNCL-ΔRGG followed by fluctuation analysis to determine the rates of recombination. All graphs are showing the rates of recombination (×10^−8^). Error bars indicate 95% confidence intervals.

To confirm the above results, we repeated the experiments by generating a series of C-terminal truncation mutants at the endogenous *NSR1* locus. As shown in Figure S2B, the protein expression levels were similar for the full length Nsr1, Nsr1ΔRGG and Nsr1ΔRBD2ΔRGG as detected by anti-Nsr1 or anti-FLAG antibody. We could not detect protein for the *nsr1ΔRBD1,2ΔRGG* allele. We confirmed that the 3XFLAG tag at the C-terminus does not interfere with the function of Nsr1 as the recombination rate of a *top1*Δ *nsr1-3XFLAG* strain was similar to the *top1*Δ strain (Figure S2C). The rates of recombination at the *pTET-lys2-GTOP* cassette in cells expressing the Nsr1ΔRBD1,2ΔRGG, Nsr1ΔRBD2ΔRGG and Nsr1ΔRGG were similar to that in cells with the complete deletion of Nsr1 (*nsr1*Δ). These results further support the requirement for the RGG domain of Nsr1 in promoting G4-associated instability.

The deletion of *NSR1* gene (*nsr1*Δ) leads to a marked slow-growth phenotype [46, 47]. We measured cell growth and calculated doubling time in *WT*, *top1Δ, nsr1Δ, top1Δ nsr1Δ,* and *top1*Δ *nsr1ΔRGG* strains. While the doubling time of a *WT* (~96minutes) and *top1*Δ strain (~99 minutes) was comparable, the *nsr1*Δ mutant strain showed significantly longer doubling time of 141 minutes (Figure 3D). This growth defect was slightly further aggravated upon the deletion of *TOP1* gene; the doubling time was ~152 minutes for the *top1*Δ *nsr1*Δ mutants. However, there was no apparent growth defect for *top1*Δ *nsr1ΔRGG* with its doubling time being similar to *WT* or *top1*Δ strain. Nsr1 has been reported to be involved in pre-rRNA processing in yeast and the deletion of *NSR1* gene leads to the defective 35S pre-rRNA processing; 35S accumulates and 20S is greatly reduced [47]. We carried out Northern hybridization analysis using oligonucleotide probes that were designed to hybridize to pre-rRNA and confirmed the sharp decrease in the 20S and the concomitant accumulation of unprocessed 35S in our *nsr1*Δ strain (Figure 3E and 3F). The deletion of *TOP1* gene had no or little effect on the pre-rRNA processing in *WT* or *nsr1*Δ backgrounds, respectively. However, in the *top1*Δ *nsr1ΔRGG* strain, the pre-rRNA processing efficiency was similar to that of the *WT* strain with a similar ratio of 20S/35S. We also observed no differences in 7S pre-rRNA level within the different mutants (Figure 3E). Overall, the above results suggest that, although the RGG domain plays an essential role in the elevation of G4-associated recombination, it is dispensable for the normal cell growth and pre-rRNA processing function of Nsr1.

When we expressed the full length Nsr1 and Nsr1ΔRGG in *WT* strains with the normal endogenous level of Nsr1, there was no effect of overexpression of the full length Nsr1 constructs while the Nsr1ΔRGG construct reduced the rates of recombination slightly (Figure 4A). In the *top1*Δ strain, the overexpression of full length Nsr1 elevated the rates of recombination at the *pTET-lys2-GTOP* cassette by ~4.5-fold compared to vector alone (Figure 4B), consistent with the 4-fold higher Nsr1 protein levels in the *top1*Δ strain overexpressing Nsr1 (Figure S2D). When the Nsr1ΔRGG construct was expressed in the *top1*Δ strain, the rates of recombination were consistently lower than the empty vector (Figure 4B). The N-term Nsr1 (1-171) was unable to induce recombination and showed rates of recombination that were similar to the empty vector. The C-term Nsr1 (172 - 414) expression resulted in about 2-fold higher recombination rates (Figure 4C). These results suggest that Nsr1 increases recombination in a dose dependent manner, and this function requires the RGG domain.

We also tested whether the highly conserved human nucleolin (NCL) can complement the loss of Nsr1 in yeast. When NCL was expressed from a plasmid, the rates of recombination at the *pTET-lys2-GTOP* cassette was elevated by ~2-fold (Figure 4D and S2E). And like the yeast Nsr1ΔRGG, the human NCLΔRGG with C-terminal deletion failed to increase the rate of recombination in *top1*Δ *nsr1*Δ cells. In *top1*Δ cells, the expression of NCL, but not NCLΔRGG resulted in further 2-fold increase in the recombination at the *pTET-lys2-GTOP* indicating that the function of Nsr1 with intact RGG domain in mediating the G4-associated genome instability is conserved in the human homolog (Figure 4D).

### Nsr1 binds to the G4 DNA *in vitro* and *in vivo*

In order to test whether the binding of Nsr1 to G4 DNA is important for the observed effect of Nsr1 in the *pTET-lys2-GTOP* and *–GBTM* constructs, we performed a gel-shift assay using the Cy5 labeled 45-mer oligonucleotide partially representing the Sμ-G sequence (Sμ-G), which was previously shown to fold into G4 structures [34], and a non-G4 forming single stranded (SS) DNA as a control (Figure 5A). G4 DNA formation by the Sμ-G oligo was confirmed by its binding to the G4-specific recombinant antibody BG4 in the presence of the G4-stabilizing K+ cation [54]. Whole cell extracts from *top1Δ, top1Δ nsr1-3XFLAG,* and *top1*Δ *nsr1*Δ cells were incubated with the folded Sμ-G oligo with or without 100-fold excess of the unlabeled competitor Sμ-G or SS DNA. Figure 5B shows a specific complex of protein bound to the Sμ-G probe in the presence of 100 mM KCl, which disappeared in the presence of the cold unlabeled Sμ-G4 competitor (band indicated by the red arrow). This band was not affected by the unlabeled SS DNA competitor and was absent in the *top1*Δ *nsr1*Δ control cells. Sμ-G oligo folded in Li+ showed a different pattern of bands when incubated with WCE from *top1*Δ cells (Figure S3A). There is a band with similar mobility as the arrow-pointed band in Figure 5B but this band (*) was competed out by the cold Sμ-G and SS-DNA indicating that this complex is not dependent on the G4 DNA conformation.

**Figure 5.**
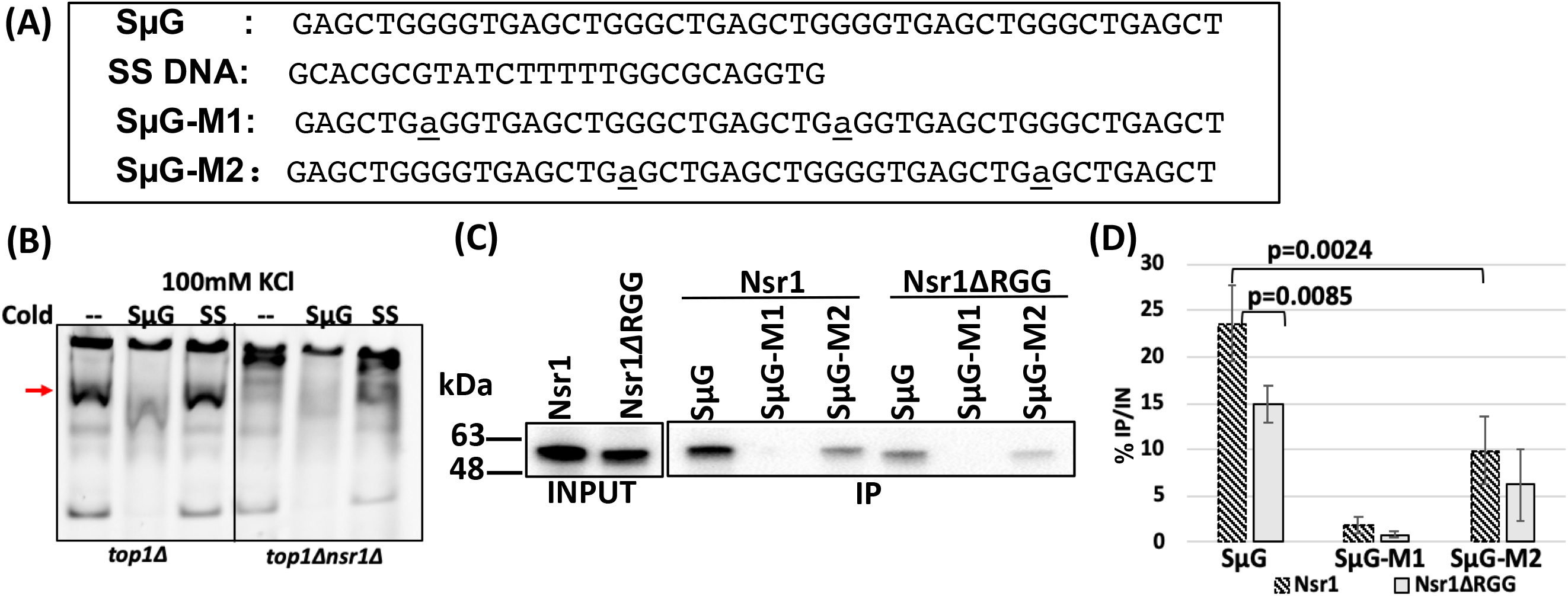
Nsr1 binds to G4 DNA *in vitro*. (**A**) Sequence of the oligos used in EMSA and pulldown assays. **(B)** Electrophoretic mobility shift assay. Cy5-conjugated Sμ-G was incubated with WCE prepared from *top1*Δ or *top1*Δ *nsr1*Δ cells, in the presence of 100mM KCl. Where indicated, a cold competitor oligo were added in 100-fold excess of the Cy5-Sμ-G. **(C)** For pull-down assays, the indicated, biotin-conjugated oligos were incubated with total yeast lysate from *top1*Δ cells expressing the C-terminal 3XFLAG tagged version of full length Nsr1, Nsr1ΔRGG, or Nsr1ΔRBD2,ΔRGG. αFLAG antibody was used to detect the respective proteins. **(D)** Quantification of % IP to input from (C); N=3.

Next, we performed affinity purification using biotinylated oligos (Figure 5A). After the denaturing-slow cooling step, each of the oligos were incubated with the whole cell lysates and pulled down using the streptavidin-coated beads. When the oligo-bound proteins were resolved by Western blot, Nsr1 was pulled downed by the G4-forming Sμ-G oligo. Nsr1-binding was significantly reduced when the runs of guanines were disrupted by G->A mutations in the second and the fourth guanine-runs (SμG-M2) and completely abrogated by G->A mutations in the first and the third guanine-runs (SμG-M1) (Figure 5C and 5D). For the Nsr1ΔRGG mutant, the binding to Sμ-G was significantly weaker compared to the full length Nsr1. The full length Nsr1, Nsr1ΔRBD2ΔRGG, and Nsr1ΔRGG were also pulled down by the anti-NCL aptamer AS1411 (GGTGGTGGTGGTTGTGGTGGTGGTGG) (Figure S3B) [55].

To determine whether Nsr1 interacts with G4 structures *in vivo*, we measured the association of Nsr1 and G4 by using chromatin immuno-precipitation (ChIP) assay. ChIP was performed in *WT* and *top1*Δ cells containing either the *pTET-lys2-GTOP* or *–GBTM* construct and expressing Nsr1 with a C-terminal 3XFlag tag. Following the pull-down with αFlag antibody-conjugated beads, qPCR analysis was done to determine the enrichment of Nsr1 at a locus 100 bp from the G4 insertion site (“G4 insert”) and a locus ~3 kb away from the G4 insertion site (“3 Kb away”). We observed ~2-fold enrichment of Nsr1 at the G4 locus when the G4 sequence was in the *GTOP* orientation, but not in the *GBTM* orientation in a *top1*Δ background (p<0.0001) (Figure 6A and 6B). Further, the enrichment of Nsr1 was significantly higher at the switch region G4 sequence insertion site than the 3’ region of the *lys2* sequence 3 kb away (p=0.0021) (Figure 6A). Nsr1ΔRGG, which is missing the C-terminal RGG domain, however, was not significantly enriched at the G4 locus in *top1*Δ cells (p=0.0009 compared to full length Nsr1) (Figure 6A). These data indicate that Nsr1 specifically associates with the G4 DNA accumulated under high transcription conditions and that this association between G4 and Nsr1 requires the C-terminal RGG repeats.

**Figure 6.**
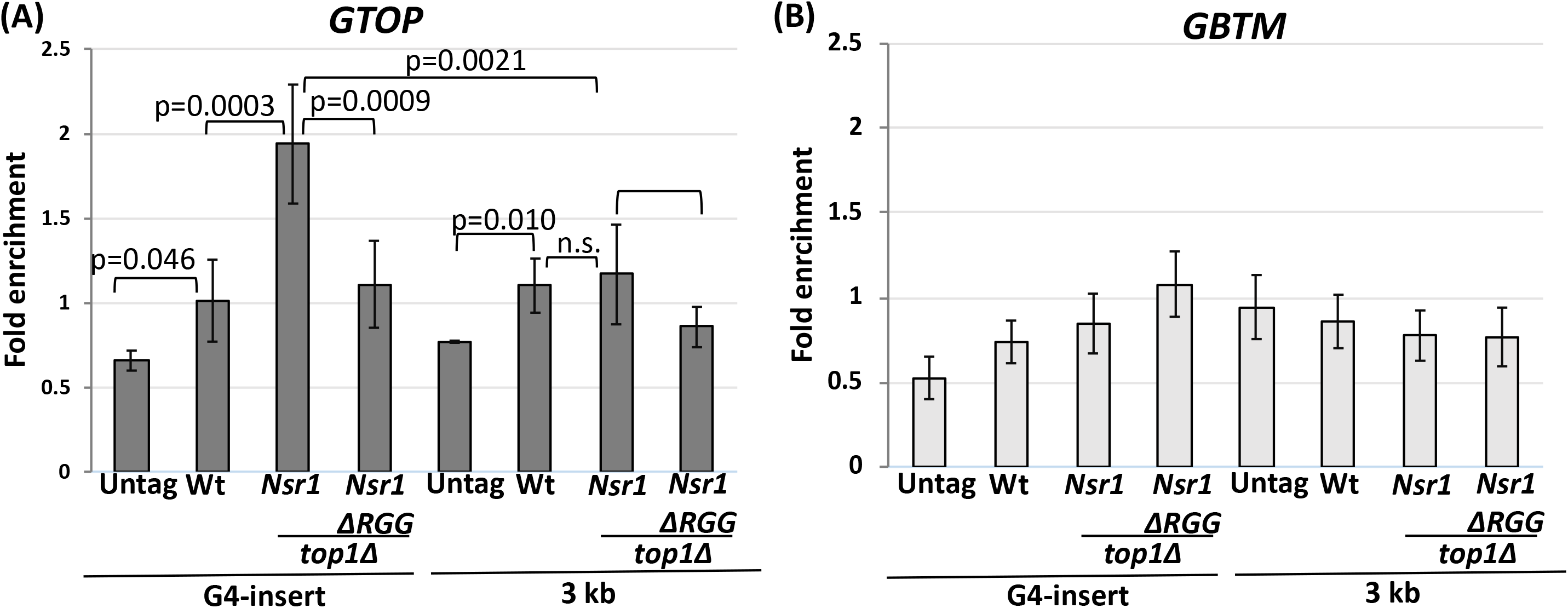
Nsr1 binding to G4 DNA *in vivo* requires the RGG domain. ChIP was carried out with an αFlag antibody and chromatin fractions prepared from yeast cells with indicated genetic backgrounds. 5΄BGL and STP primers were previously described and anneal within the *LYS2* ORF at ∼100 and ∼3000 bp from the Sμ sequence insertion site, respectively. All values are based on at least six independent samples with the exception of the no-tag samples (N=3) and the standard deviations are indicated by error bars. P-values shown were calculated using student t-test. **(A)** GTOP strains **(B)** GBTM strains.

### Nsr1-disruption allows for the better access by activation-induced deaminase to the single stranded DNA in the G4-prone sequence

Activation-induced deaminase (AID) converts cytosine to uracil and is required for the somatic hypermutation (SHM) of immunoglobulin variable region genes as well as the heavy chain class switch recombination (CSR) [56]. We previously showed that the ectopic expression of human AID in the *WT* and *top1*Δ yeast cells resulted in a higher rate of recombination at the *pTET-lys2-GBTM* than *-GTOP* in both strain backgrounds [21]. This difference was partly attributed to the larger number of cytosines present on the single-stranded non-transcribed strand (NTS) in the *GBTM* orientation compared to the *GTOP* orientation (358 vs. 117 cytosines) since AID specifically targets the cytosines located on the single-stranded DNA. We postulated that NSR1-G4 DNA complex could further impede the accessibility of AID to those cytosines on the NTS in the *pTET-lys2-GTOP* construct located proximal to the G4 structure. Thus, we tested whether the cytosines present on the NTS in the GTOP-orientation could become more accessible to AID in the absence of Nsr1 protein. When we expressed hAID, the rate of recombination at the *pTET-lys2-GBTM* cassette was elevated by about 4- to 5-fold in the *WT*, *top1Δ,* and *top1*Δ *nsr1*Δ strains. For the *pTET-lys2-GTOP* construct, the rate of recombination was not changed by the expression of hAID in *WT* and *top1*Δ strains, but increased by about 3-fold in a *top1*Δ *nsr1*Δ strain (Table 1). This increased rate of recombination could reflect the less-restricted access of hAID to the cytosines in the absence of G4 DNA-Nsr1 complex on the NTS.

**Table 1:** Effect of hAID expression on G4-associated recombination rates.

|  | Genotype | Recombination rate x 10<sup>-8</sup> (CI*) |  | Fold Change:<br>hAID/Vector |
| --- | --- | --- | --- | --- |
|  |  | Empty vector | hAID |  |
| *pTET-lys2-*<br>*GTOP* | *WT* | 20.4 (15.1-27.9) | 31.7 (16.2-60.2) | 1.5 |
|  | *top1Δ* | 220 (177-322) | 219 (137-661) | 0.99 |
|  | *top1Δnsr1Δ* | 20.6 (13.7-27.8) | 54.1 (46.3-65.8) | 2.6 |
| *pTET-lys2-*<br>*GBTM* | *WT* | 12.7 (8.0-17.1) | 59.1 (44.7-84.5) | 4.6 |
|  | *top1Δ* | 11.4 (8.92-15.3) | 45.6 (38.8-51.4) | 4.0 |
|  | *top1Δnsr1Δ* | 10.4 (8.3-15.5) | 56.7 (29.7-150) | 5.4 |

\* 95% confidence interval

### Nsr1 obstructs replication at a G4 DNA-containing locus

The above genetic and physical data and the enrichment of Nsr1 at the G4 –insert loci suggests that Nsr1 binds tightly to G4 DNA *in vivo* and obstruct replication. In order to determine whether Nsr1-G4 DNA complex forms a replication block, we measured the DNA copy number of specific genomic loci throughout the S phase. Droplet digital PCR (ddPCR), which provides an absolute quantification of the target DNA with high precision, accuracy, and sensitivity [57], was used to determine the replication kinetics as inferred from the copy number changes throughout the S phase (see Materials and Methods). Our approach was modified from a previously reported use of ddPCR in determining the replication timing of multiple sites in the yeast and human genomes [58]. To determine the locus-specific replication timing, cells were first arrested with α-factor and released into the S phase. Flow cytometry confirmed synchronous progression through S phase upon release from α-factor (Figure S4A and S4B). The DNA samples collected at multiple time points from 0 to 100 minutes after the release were used to determine the copy numbers at three different loci; “ARS306” – near the early-firing autonomously replicating sequence on chromosome III, “KanMX” - between “ARS306” (~8 Kb distal) and the *pTET-lys2-GTOP* cassette (~2 Kb distal), and “STE50” – further distal from “ARS306” (~14 Kb distal) compared to the *pTET-lys2-GTOP* cassette (~4 Kb distal) (Figure 7A). The other nearest origin of replication is *ARS305,* which is located >23 Kb from the “STE50” locus. The replication in this genomic region was previously determined to predominantly originate from *ARS306* [59]. The replication kinetics at these three genomic loci in yeast cells in a *top1*Δ background with either *NSR1* or *nsr1ΔRGG* allele were compared; the significant growth defect in *nsr1*Δ cells compared to *NSR1* cells, which is not present in cells with the *nsr1ΔRGG* allele (Figure 3D), would have interfered with a straightforward comparison. In respect to the recombination occurring at the *pTET-lys2-GTOP* cassette, the *nsr1ΔRGG* allele is indistinguishable from the *nsr1*Δ null allele (Figure 3B and 4B).

**Figure 7.**
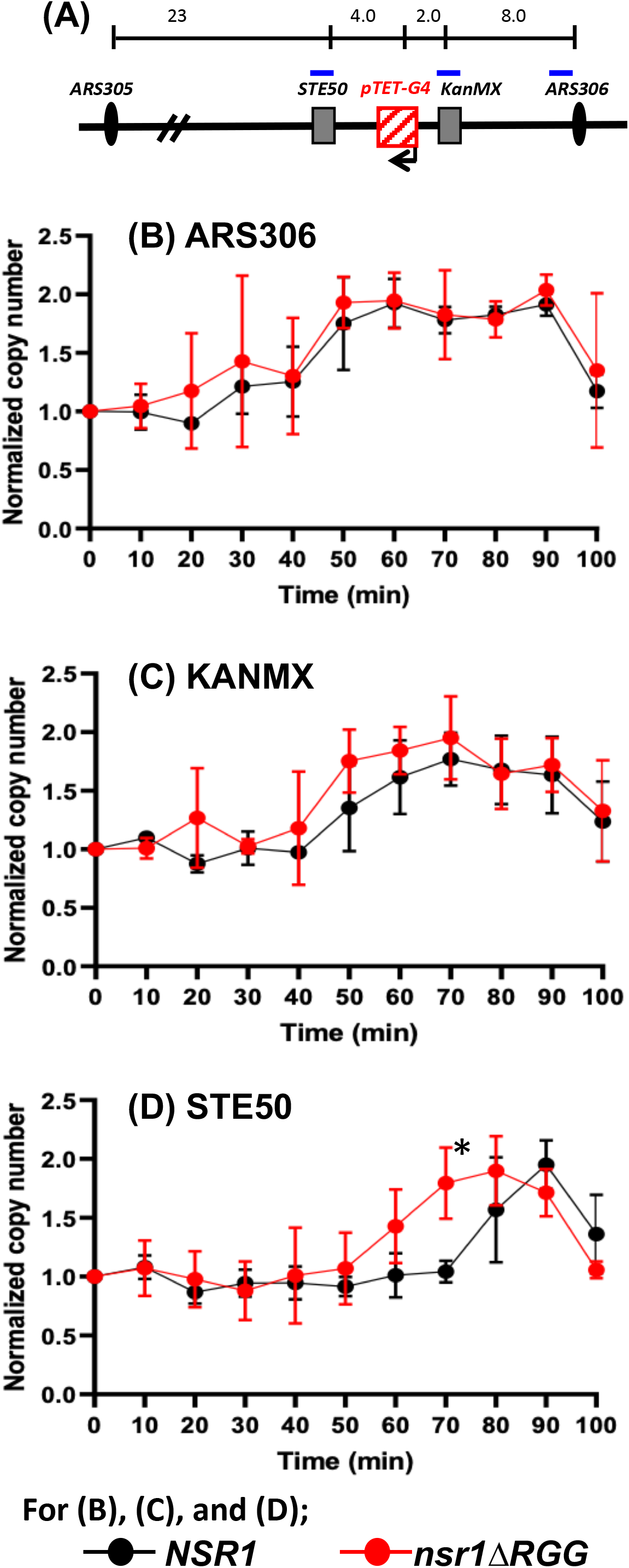
DNA replication timings and copy number changes determined by ddPCR. (**A**) A schematic drawing of the regions proximal to the *pTET-lys2-GTOP* cassette on yeast chromosome III. The locations of primers used in ddPCR are indicated by blue bars. For the sequences of the primers, see Table S1. The distance indicated above in Kb are estimates and not to scale. (**B - D**) DNA samples analyzed by ddPCR were extracted at the indicated time points (minutes after the release from α-factor). For calculating the copy number, each time point values of “ARS306”, “KanMX” and “STE50” loci were normalized to time 0 value of the “ARS306” locus. And for “KanMX” and “STE50” loci, they were further normalized to the “KanMX” at time 0 and “STE50” at time 0, respectively. Data from at least 3 independent experiments was used to calculate mean and standard deviations (indicated by error bars). Normalized copy number of (**B**) “ARS306” (**C**) “KanMX” and (**D**) “STE50” loci. P-values was calculated using student t-test. Asterisks denote P-value =0.0149.

During the S phase, the copy number at each genomic locus is expected to progressively increase from one to two as replication proceeds. The time at which replication completes and the copy number reaches two will be mostly dependent on the proximity to the relevant replication origin, *ARS306*. In *top1*Δ *nsr1ΔRGG* cells containing the *pTET-lys2-GTOP* cassette, the copy number of “ARS306” locus reached two at around 50 - 60 minutes after the release from the α-factor (Figure 7B). A similar pattern for this locus was observed in *top1*Δ cells, which express the full length Nsr1 protein. For the *top1*Δ and *top1*Δ *nsr1ΔRGG* strains containing the *pTET-lys2-GBTM* cassette, the copy number of “ARS306” locus also reached two at around 50 - 60 minutes after the release from the α-factor with no significant difference between the two strain backgrounds (Figure S4C). For the “KanMX” locus in the strains with the *pTET-lys2-GBTM* cassette, the time after the release from the α-factor required to reach the copy number of two was between 60 and 70 minutes regardless of the strain background (Figure S4D). However, in the *top1*Δ cells with the *pTET-lys2-GTOP* cassette, there was a slight but statistically non-significant lag in the replication at the “KanMX” locus compared to the *top1nsr1ΔRGG* cells (Figure 7C). At the “STE50” loci located downstream of the *pTET-lys2-GTOP* cassette relative to *ARS306,* there was a more significant difference in the replication kinetics between the *top1*Δ *nsr1ΔRGG* and *top1*Δ strains (Figure 7D). While the copy number of this locus reached two around 70 – 80 minutes after the release from the α-factor in *top1nsr1ΔRGG* cells, in the *top1*Δ cells, the copy number of “STE50” was significantly below two at the 70-minute time point, reaching two only 90 minutes after the release from the α-factor. For the *top1*Δ and *top1*Δ *nsr1ΔRGG* strains containing the *pTET-lys2-GBTM* cassette, no difference in the replication kinetics was observed with the copy number at “STE50” reaching two around 70 – 80 minutes after the release from the α-factor (Figure S4E).

## DISCUSSION

Although initially described as a sequence-specific RNA-binding protein, human nucleolin (NCL) preferentially binds to G-quadruplex structures over RNA substrates and plays a crucial role in G4 metabolism [17, 39, 40]. In this study, we have explored the possibility that the yeast homolog Nsr1 also contributes to G4 DNA metabolism. As expected from its highly conserved RBD and RGG domains, we demonstrated that the physical association with G4 DNA is a conserved function in Nsr1. Both G4-forming oligos (Sμ-G and AS1411) and non-G4-forming oligos (SS DNA, Sμ-G M1, and Sμ-G M2) were used to show that Nsr1 is a specific G4 DNA binding protein (Figure 5B, 5C, and S3). Nsr1 is also significantly enriched *in vivo* at the G4 reporter construct only when the guanines are on the top, non-transcribed strand of an actively transcribed gene (i.e. *pTET-lys2-GTOP*) in a *top1*Δ background, which is the condition with the significant elevation of G4-associated recombination (Figure 6A). Nsr1-enrichment is significantly reduced ∼3 kb away from the G4 motif sequence even though it is within the same transcribed unit, which further supports the specificity of Nsr1 interaction with G4 DNA. Although the *in vitro* pull-down assay showed that Nsr1ΔRGG can bind G4 DNA forming oligos, ChIP experiments showed that *in vivo* binding of Nsr1 to a G4 DNA-containing genomic locus is dependent on the presence of the RGG domain.

The disruption of Nsr1 substantially decreases the G4-associated genomic instability as manifested by the decreased rates of recombination and of GCR in *top1*Δ background by about 4- and 7-fold, respectively (Figures 1 and 2). The reduction in genome instability in the absence of Nsr1 was not specific to the switch region G-rich sequence as GCRs occurring at a different G-rich motif from the human TCF3 gene were also reduced upon deletion of *NSR1* (Figure 2D). Additionally, overexpression of Nsr1 led to further elevation of recombination specifically at the *pTET-lys2-GTOP* cassette in a *top1*Δ background (Figure 4B). One interpretation of these surprising results is that the consequence of the association between Nsr1 and G4 DNA is the elevated genome instability. The genome instability associated with G4-structures should be prevented by the activity of G4-unwinding DNA helicases including yeast Sgs1 and Pif1 [60, 61]. We considered the possibility that Nsr1 can occlude DNA helicases from recognizing and then resolving G4 DNA. Human nucleolin NCL, in fact, was reported to prevent Werner helicase from unwinding G4 oligos in *vitro* [62]. When we expressed hAID to induce the cytosine deamination and subsequent recombination, those cytosines present on the NTS at the G4 DNA-forming *pTET-lys2-GTOP* cassette were more susceptible to hAID processing in a *top1*Δ *nsr1*Δ background than in a *top1*Δ background, supporting the idea that the regions proximal to the G4 DNA are in a less accessible conformation in the presence of Nsr1.

In the mammalian systems, NCL was shown to localize to the sites of DNA double strand breaks (DSB) and to reorganize nucleosomes proximal to DSBs [63]. Since this role of NCL is critical in the early step of DSB repair by homologous recombination (HR), there is a possibility that the reduction in the recombination rate at the *pTET-lys2-GTOP* cassette in *nsr1*Δ yeast cells is due to the general defect in HR. This alternative explanation for the role of Nsr1 is contradicted by the fact that the *NSR1*-deletion did not affect the recombination rate of the *pTET-lys2-GBTM* construct in *top1*Δ background or the recombination rates of either constructs in a *rnh1Δ rnh201*Δ background (Figure 1B and 1D). Also, the deletion of *NSR1* led to the reduction in GCR events (Figure 2), which normally occur through the HR-independent, *de novo* telomere addition in haploid yeast cells [64]. Finally, in order to rule out any defect in DNA damage checkpoint response, we checked the phosphorylation state of Rad53 in the *WT*, *top1*Δ, *nsr1Δ,* and *top1Δnsr1*Δ cells treated with the DSB-inducing drug Zeocin. After 90 mins with Zeocin, Rad53 phosphorylation was apparent in all cell types indicating that the DNA damage checkpoint response was not affected by the disruption of Nsr1 (Figure S4F).

Located C-terminal to the multiple RNA binding domains (RBDs), yeast Nsr1 and human NCL contain 6 and 9 repeats of arginine-glycine-glycine or RGG motifs, respectively, which are found in proteins associated with important nucleic acid-mediated functions such as transcription, pre-mRNA splicing, DNA damage signaling and mRNA translation [65]. Other proteins with verified roles in DNA metabolism and cancer development that contain RGG motifs include Mre11 (a DSB processing enzyme), Mll4 (a histone methyltransferase), and Ews (Ewings Sarcoma protein; DNA damage response protein). Recently, the RGG-box of hnRNPA1, a member of ribonucleoproteins, was shown to bind specifically to the telomeric G4 DNA [44]. In yeast Nsr1, RGG motifs also appear to play an important function. Yeast cells expressing Nsr1 with the truncation of this motif (Nsr1ΔRGG) does not show the severe defects in the pre-rRNA processing and cell growth that are typical of the *nsr1*Δ cells (Figure 3D - 3F). However, when we deleted the RGG domain from Nsr1, the resulting Nsr1ΔRGG construct lost the ability to complement the full-length Nsr1 in promoting G4-associated genome instability at the *pTET-lys2-GTOP* cassette either in *top1*Δ *nsr1*Δ or *top1*Δ background (Figure 3B and 4B). This can be due to weaker interaction between G4 DNA and the Nsr1ΔRGG mutant protein. Earlier biochemical analysis showed that the RGG domain of hNCL is required for high-affinity interaction with G4-forming oligos in *vitro* [45]. We showed that the in *vitro* interaction between the yeast Nsr1 and G4 oligos Sμ-G as well as the genomic enrichment of Nsr1 protein to the co-transcriptionally formed G4 DNA become significantly weaker when the RGG domain is deleted (Figure 5C, 5D, and 6A).

As our data indicates, the elevation of G4-associated genomic instability by Nsr1 requires a strong physical interaction with G4 DNA as mediated by the RGG domain. Such conclusion suggests that Nsr1 binding leads to stabilization of the co-transcriptionally formed G4 DNA and that the Nsr1/G4 DNA forms a nucleoprotein complex that acts as a replication block. Non-histone protein-DNA complexes forming a replication block have been previously reported for the origin of replication (ORC) complex and Rap1 [66]. In order to determine whether the Nsr1-G4 DNA complex poses a physical replication obstruction, we measured the replication kinetics of the genomic region proximal to the G4 DNA-containing *pTET-lys2-GTOP* cassette using the ddPCR approach, where the copy number of the specific genomic loci were tracked through a single S phase. As shown in Figure 7A, replication proceeds from *ARS306* through “KanMX” regions and the *pTET-lys2-GTOP* cassette before it passes “STE50” locus. As would be expected from the replication block forming at the *pTET-lys2-GTOP*, no significant deviation in the replication kinetics was observed at the sites located between the replication origin and the *pTET-lys2-GTOP* (i.e. “ARS306” and “KanMX” Figure 7B and 7C). However, the replication fork must process past the G4-containing *pTET-lys2-GTOP* region before it reaches “STE50”, where the significant delay in replication was observed in the presence of Nsr1 in *top1*Δ background (Figure 7D). This delay in replication progress, just as the elevated G4-associated recombination, required the RGG domain of Nsr1; replication at “STE50” proceeded with significantly faster kinetics in cells expressing Nsr1ΔRGG compared to cells expressing the full length Nsr1. This difference between Nsr1ΔRGG and Nsr1 further supports the idea that the strong physical interaction between Nsr1 and G4 DNA underlies both G4-specific replication obstruction and genome instability. In further confirmation that the replication obstruction by Nsr1 is G4-specific, in cells containing the non-G4 forming *pTET-lys2-GBTM* cassette, the replication kinetics at all three loci showed no significant differences whether in *NSR1* or *nsr1ΔRGG* background. Even the “STE50” locus replicated with similar relative replication timings (Figure S4E).

In summary, we have identified a novel function of yeast Nsr1 as a G4 DNA binding protein. Nsr1 shows strong *in vitro* binding to G4 forming sequences and is specifically enriched at a co-transcriptionally formed G4 DNA *in vivo*. Moreover, we first demonstrate here that Nsr1 interaction with G4 DNA results in very significant replication impediment in a G4 DNA-specific manner. This key finding suggests that the formation of a stable Nsr1-G4 DNA complex functioning as a replication obstruction underlies the significant elevation in G4-associated genome instability. Importantly, the data presented here point to the requirement of the conserved RGG domain at the C-terminal end of Nsr1 in promoting the instability at G4 DNA. This result calls for further study into the conformational changes associated with the RGG domain in the Nsr1-G4 DNA complex that could enable such a complex to function as a replication obstacle.

## MATERIALS AND METHODS

### Yeast strains and plasmids

Yeast strains used for the mutation and recombination assays were derived from YPH45 (*MATa, ura3-52 ade2-101 trp1Δ1*). Construction of strains containing the *pTET-lys2-GTOP or – GBTM* constructs were previously described [21]. Gene deletions were carried out via the one-step allele replacement by amplification of loxP-flanked marker cassettes. Nsr1-expression plasmid was constructed by amplifying NSR1 ORF along with 490 nt upstream and 250 nt downstream from the yeast genomic DNA and cloning into the yeast *CEN* vector pRS316. The deletion constructs, N-Term Nsr1 and C-term Nsr1, have been previously described [67]. The expression plasmid for human Nucleolin was constructed by amplifying the NCL ORF from GFP-Nucleolin (Addgene; #28176) using primers NCL-For and NCL-1XHA Rev (Table S1). The BamHI/ XhoI digested NCL PCR product was cloned into BamHI/ XhoI digested pGPD2 (Addgene; #43972). The NCL-ΔRGG plasmid was similarly created using primers NCL-For and NCLΔRGG-1XHA Rev (Table S1). The expression of NCL and NCL-ΔRGG was checked by anti HA-HRP antibody (Sigma; H6533).

### Determination of rates

Recombination rates and 95% confidence intervals were determined using the method of the median as described previously [22]. Twelve to twenty-four individual cultures were used to determine each rate and the associated 95% confidence intervals. Recombination rates are considered to be statistically different when the 95% confidence intervals, which are indicated in each graph as error bars, do not overlap. For the gross chromosomal rearrangement (GCR) assay, 5 ml cultures in YEPD medium (1% yeast extract, 2% Bacto-peptone, 2% dextrose, and 250 μg/mL adenine hemisulfate) were inoculated with single colonies and grown for 3 days at 30°C. Cells were then plated either on YEPD-agar or synthetic complete dextrose medium lacking arginine (SCD-arg) and containing canavanine (60 mg/L) and 5-Fluoroorotic acid (5-FOA; 1 g/L) [52].

### Western blotting

Whole cell extracts (WCE) prepared using NaOH lysis method were separated on a 4-20% polyacrylamide gel (BioRad), and transferred using a Semi-Dry Trans-Blot Cell (BioRad). After blocking, anti-FLAG (Sigma, A8592) and anti-GAPDH (Invitrogen, MA5-15738) antibodies were used for detection. The blot was developed using West-Q Pico Dura ECL Solution (GenDepot) and the ChemiDoc MP Imaging system (BioRad). Rad53 phosphorylation was checked using Rad53 antibody (ab104232) from Abcam.

### Growth curve

Growth curve and doubling time measurement: eight independent cultures of each genotype were grown to mid-logarithmic phase, diluted to ~1 × 10^6^ cells/ml and incubated in 96-well plate at 30°C. OD_600_ was measured automatically every 10 min. Doubling time was calculated using the following equation: Doubling time = [hours cells grown Ln(2)]/[Ln (Nt / N0)], where Nt and N0 are the OD_600_ at two different times in log phase of growth.

### Northern blot analysis

Northern blot analysis of the ribosomal RNAs was carried out as previously described [68].

### Electrophoretic mobility shift assays (EMSA)

Yeast extract preparation was carried out according to a previously described protocol [69]. For binding assays, 5΄ Cy5 labeled oligos (Sigma) were resuspended in either 100 mM KCl/10 mM Tris–Cl or 100 mM LiCl/10 mM Tris–Cl, denatured at 95°C for 5 min and finally slowly cooled overnight to allow folding. Unlabeled, cold competitor oligos were treated in parallel to the labeled oligos. For binding reactions, approximately 10–15 μg of protein from each cell lysate was incubated with annealed oligos (0.2 μM) in 100 mM KCl or LiCl, 20 mM HEPES pH 7.5, 0.01% NP40, 5% glycerol, 5 mM MgCl_2_ and incubated at RT for 15 min before running on 15% non-denaturing TBE-polyacrylamide gel. Gel images were captured using BioRad Chemidoc imager.

### DNA-Dynabeads Affinity Purification of Proteins

The DNA-Dynabeads affinity purification of proteins was carried out according to the previously described protocol [70] with some modifications. For DNA-conjugated Dynabeads preparation, biotinylated oligonucleotides SμG, SμG-M1, SμG-M2, and AS1411 (Sigma) were incubated at 60°C overnight in the presence of 10 mM Tris pH 7.5 and 100 mM KCl and then conjugated to Streptavidin-Coupled M-280 Dynabeads® (Life Technologies) as per the manufacturer’s instructions. Yeast cells from indicated backgrounds were used to prepare whole cell extracts by glass bead-mediated cell disruption. The oligo-conjugated beads were incubated with the yeast extract at 4 °C. After washing, proteins were eluted by boiling in 1XSDS-PAGE loading buffer followed by immunoblotting analysis using anti-FLAG antiboy (Sigma; Cat# A8592).

### Chromatin immunoprecipitation

For Chromatin IP (ChIP), a previously described protocol was used [34] with following modifications. Anti-FLAG antibody conjugated beads (Life Technologies) were used in pull-down. Ct values for each ChIP samples were first normalized to the corresponding input samples, and then divided by the values for the *CAN1* locus to calculate the relative fold enrichment. Primers used in the qPCR analysis were previously described [34]. P-values were calculated using the student t-test.

### Cell synchronization and time course experiment

Cell synchronization and sample collection was carried out as described with slight modifications [71]. Briefly, 30°C grown, log phase cells (OD_600_~0.5 - 0.6) with *bar1*Δ were arrested in late G1 phase using 50 ng/ml alpha-factor peptide (Sigma) for ~120 minutes, released by washing with fresh YEPD medium with 50μg/ml Pronase (Sigma-Aldrich), and then allowed to proceed through a synchronous cell cycle at 21°C. Samples were removed at 10 min intervals and immediately treated with buffer containing 0.1% sodium azide. For flow cytometry, yeast cells were fixed overnight in the ice-cold 70% ethanol at 4°C fixed and then re-suspended in 500μl of 50μmM sodium citrate (pH 7.2) containing RNase A and incubated for 2 hours at 37°C, followed by incubation with 50μl of 20μmg/ml Proteinase K. at 50°C for 1 - 2 hours. Cells were stained with SYTOX Green dye (4μμM in 50mM sodium citrate, pH 7.2; Thermo Scientific) for at least one hour and then analyzed using the BD FACS ARIA II SORP and FlowJo software (FlowJo, LLC).

### DNA extraction and ddPCR

Yeast genomic DNA extraction from the cells collected at 10-minute intervals after the release from alpha-factor was carried out as described [58]. DNA concentration was measured using the Qubit™ dsDNA HS assay (ThermoFisher). The DNA was sonicated for 2 cycles of 15sec ON/90 sec OFF in a bioruptor (Diagenode) at 4°C. The ddPCR reaction consisted of 1.5–2 ng of sonicated genomic DNA, 1× QX200 ddPCR EvaGreen Supermix (Bio-Rad) and primers (200 nM final each). The samples were processed using the QX200 Droplet Digital PCR system and analyzed with the QuantaSoft software (Bio-Rad). The primers are listed in Supplementary table S1. For calculating the copy number, each time point values of the “ARS306”, “KanMX”, and “STE50” loci were normalized to time 0 value of the “ARS306” locus and for the “KanMX” and “STE50” loci, they were further normalized to the “KanMX” (time 0) and “STE50” (time 0), respectively. Data from at least 3 independent experiments was used to calculate the standard deviation. p-values were calculated by using student t-test (Graphpad Prism).

## Acknowledgements

Authors would like to thank Dr. A. Saiardi of University College London, London UK for the gift of plasmids with the C-term and N-term Nsr1 constructs. We also thank the members of Kim lab for discussion and critical reading of the manuscript.

## Supplementary Figure Legends

**Figure S1.** Rates of recombination of the indicated strains under low transcription conditions.

**Figure S2.** RGG domain of Nsr1 is required to induce high recombination in *top1*Δ background. Whole cell extracts (WCE) from the indicated yeast strains were prepared to check the expression of Nsr1 protein in various deletion constructs **(A)** Full length Nsr1 and Nsr1ΔRGG are expressed from a plasmid in *top1Δnsr1*Δ background and level of protein expression checked by αNsr1 antibody. **(B)** The indicated C-terminal truncations in the Nsr1 gene were generated at the endogenous loci and protein expression was detected by αFLAG antibody in the upper panel and αNsr1 antibody in the middle panel. C-terminal 3XFLAG tags were indicated by *. GAPDH in the lower panel serves as a loading control. Bars to the left indicate positions of the molecular weight markers (kDa). **(C)** The graph shows the rates of recombination (×10^−8^) of the strains mentioned in B. Error bars indicate 95% confidence intervals. Relevant genotypes are indicated. **(D)** To check the level of Nsr1 overexpression, empty vector and Nsr1 containing plasmid are expressed in *top1*Δ background and probed with αNsr1. **(E)** WCE prepared from *top1Δnsr1*Δ cells transformed with empty vector, full length human Nucleolin (NCL) and NCLΔRGG was probed with αHA antibody to check protein expression.

**Figure S3.** Electrophoretic mobility shift assay. **(A)** Cy5-conjugated Sμ-G oligos incubated with WCE prepared from *top1*Δ or *top1*Δ *nsr1*Δ cells, in buffer containing 100 mM LiCl. Where indicated, a cold competitor oligo in 100-fold excess of the Cy5-Sμ-G oligo was added. **(B)** Pull down of yeast lysate from *top1*Δ, *top1*Δ *nsr1ΔRGG, and top1Δ nsr1ΔRBD2ΔRGG* cells containing endogenous 3XFLAG tag at C-terminus with folded biotinylated AS1411 oligo.

**Figure S4.** (**A**) Flow cytometry analysis of the indicated strains containing the *pTET-lys2-GTOP* cassette. t = minutes after α-factor release. (**B**) Flow cytometry analysis of the indicated strains containing the *pTET-lys2-GBTM* cassette. **(C – E)** Synchronized GBTM strains were analyzed by ddPCR as described in Figure 7. Normalized copy number of (**C**) ARS306 (**D**) KanMX (ARS+~8Kb) and **(E)** Ste50 (ARS+17Kb) loci. **(F)** DNA damage response is functional in *top1Δnsr1*Δ background. Whole cell extracts from the indicated genotypes were prepared following mock (M) or zeocin (Z) treatment of the cells for 90 minutes. Western blot analysis using anti-Rad53 antibody was done to check Rad53 phosphorylation state.

**Table S1.**
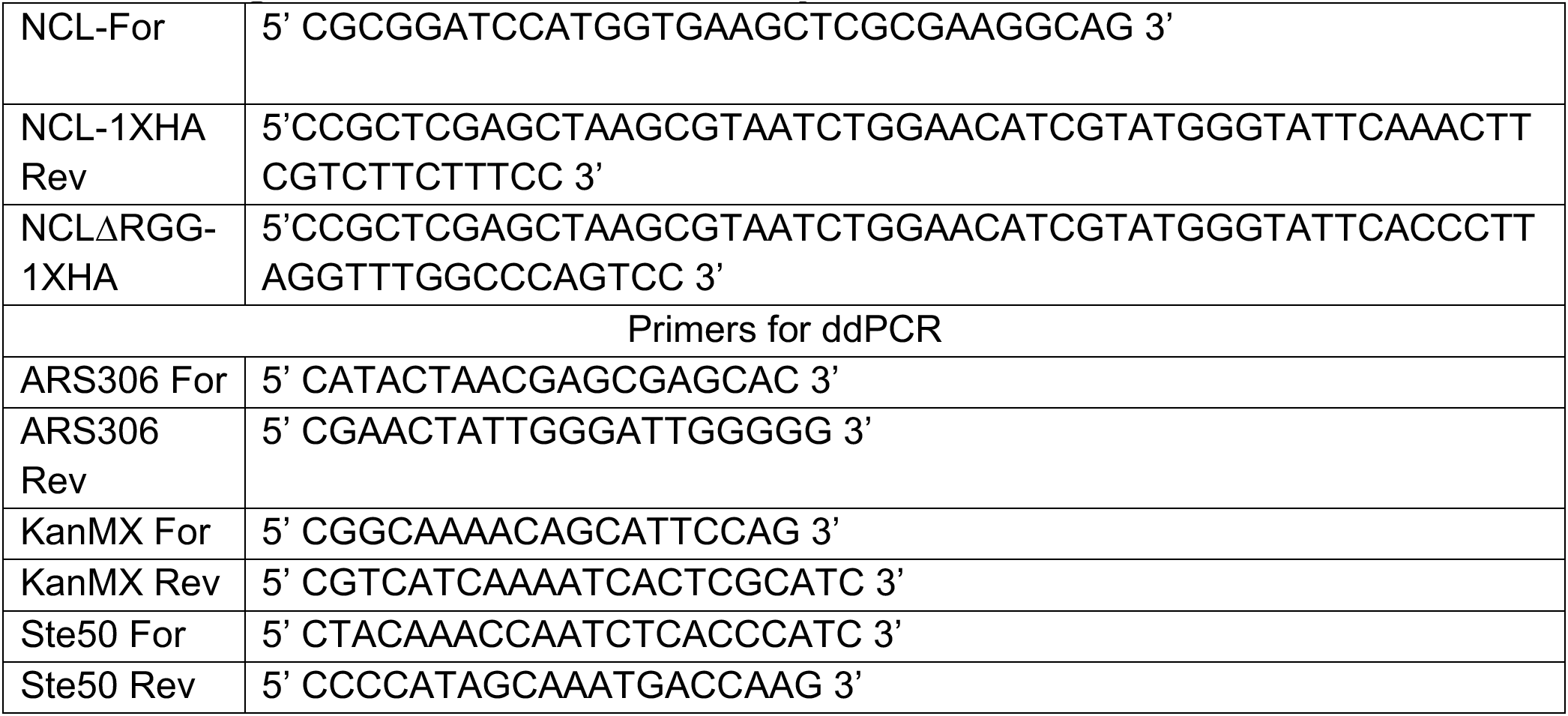
Oligonucleotides used in the study

## REFERENCES

1. Maizels N, Gray LT. The G4 genome. PLoS Genet. 2013;9(4):e1003468. Epub 2013/05/03. doi: 10.1371/journal.pgen.1003468 PGENETICS-D-13-00197 [pii]. PubMed PMID: 23637633; PubMed Central PMCID: PMC3630100.

2. Bochman ML, Paeschke K, Zakian VA. DNA secondary structures: stability and function of G-quadruplex structures. Nat Rev Genet. 2012;13(11):770–80. Epub 2012/10/04. doi: nrg3296 [pii] 10.1038/nrg3296. PubMed PMID: 23032257; PubMed Central PMCID: PMC3725559.

3. Kim N. The interplay between G-quadruplex and Transcription. Curr Med Chem. 2017. Epub 2017/12/30. doi: CMC-EPUB-87624 [pii] 10.2174/0929867325666171229132619. PubMed PMID: 29284393; PubMed Central PMCID: PMC6026074.

4. Capra JA, Paeschke K, Singh M, Zakian VA. G-quadruplex DNA sequences are evolutionarily conserved and associated with distinct genomic features in Saccharomyces cerevisiae. PLoS Comput Biol. 2010;6(7):e1000861. Epub 2010/08/03. doi: 10.1371/journal.pcbi.1000861. PubMed PMID: 20676380; PubMed Central PMCID: PMC2908698.

5. Huppert JL, Balasubramanian S. Prevalence of quadruplexes in the human genome. Nucleic Acids Res. 2005;33(9):2908–16. Epub 2005/05/26. doi: 33/9/2908 [pii] 10.1093/nar/gki609. PubMed PMID: 15914667; PubMed Central PMCID: PMC1140081.

6. Eddy J, Maizels N. Conserved elements with potential to form polymorphic G-quadruplex structures in the first intron of human genes. Nucleic Acids Res. 2008;36(4):1321–33. Epub 2008/01/12. doi: gkm1138 [pii] 10.1093/nar/gkm1138. PubMed PMID: 18187510; PubMed Central PMCID: PMC2275096.

7. Rawal P, Kummarasetti VB, Ravindran J, Kumar N, Halder K, Sharma R, et al. Genome-wide prediction of G4 DNA as regulatory motifs: role in Escherichia coli global regulation. Genome Res. 2006;16(5):644–55. Epub 2006/05/03. doi: 16/5/644 [pii] 10.1101/gr.4508806. PubMed PMID: 16651665; PubMed Central PMCID: PMC1457047.

8. Todd AK, Johnston M, Neidle S. Highly prevalent putative quadruplex sequence motifs in human DNA. Nucleic Acids Res. 2005;33(9):2901–7. Epub 2005/05/26. doi: 33/9/2901 [pii] 10.1093/nar/gki553. PubMed PMID: 15914666; PubMed Central PMCID: PMC1140077.

9. Du Z, Zhao Y, Li N. Genome-wide analysis reveals regulatory role of G4 DNA in gene transcription. Genome Res. 2008;18(2):233–41. Epub 2007/12/22. doi: gr.6905408 [pii] 10.1101/gr.6905408. PubMed PMID: 18096746; PubMed Central PMCID: PMC2203621.

10. Hershman SG, Chen Q, Lee JY, Kozak ML, Yue P, Wang LS, et al. Genomic distribution and functional analyses of potential G-quadruplex-forming sequences in Saccharomyces cerevisiae. Nucleic Acids Res. 2008;36(1):144–56. Epub 2007/11/15. doi: gkm986 [pii] 10.1093/nar/gkm986. PubMed PMID: 17999996; PubMed Central PMCID: PMC2248735.

11. Moruno-Manchon JF, Koellhoffer EC, Gopakumar J, Hambarde S, Kim N, McCullough LD, et al. The G-quadruplex DNA stabilizing drug pyridostatin promotes DNA damage and downregulates transcription of Brca1 in neurons. Aging (Albany NY). 2017;9(9):1957–70. Epub 2017/09/15. doi: 101282 [pii] 10.18632/aging.101282. PubMed PMID: 28904242; PubMed Central PMCID: PMC5636668.

12. Sarkies P, Reams C, Simpson LJ, Sale JE. Epigenetic instability due to defective replication of structured DNA. Mol Cell. 2010;40(5):703–13. Epub 2010/12/15. doi: S1097-2765(10)00847-6 [pii] 10.1016/j.molcel.2010.11.009. PubMed PMID: 21145480; PubMed Central PMCID: PMC3145961.

13. Seenisamy J, Rezler EM, Powell TJ, Tye D, Gokhale V, Joshi CS, et al. The dynamic character of the G-quadruplex element in the c-MYC promoter and modification by TMPyP4. J Am Chem Soc. 2004;126(28):8702–9. Epub 2004/07/15. doi: 10.1021/ja040022b. PubMed PMID: 15250722.

14. Siddiqui-Jain A, Grand CL, Bearss DJ, Hurley LH. Direct evidence for a G-quadruplex in a promoter region and its targeting with a small molecule to repress c-MYC transcription. Proc Natl Acad Sci U S A. 2002;99(18):11593–8. Epub 2002/08/27. doi: 10.1073/pnas.182256799 182256799 [pii]. PubMed PMID: 12195017; PubMed Central PMCID: PMC129314.

15. Bacolla A, Tainer JA, Vasquez KM, Cooper DN. Translocation and deletion breakpoints in cancer genomes are associated with potential non-B DNA-forming sequences. Nucleic Acids Res. 2016;44(12):5673–88. Epub 2016/04/17. doi: gkw261 [pii] 10.1093/nar/gkw261. PubMed PMID: 27084947; PubMed Central PMCID: PMC4937311.

16. Majounie E, Abramzon Y, Renton AE, Perry R, Bassett SS, Pletnikova O, et al. Repeat expansion in C9ORF72 in Alzheimer’s disease. N Engl J Med. 2012;366(3):283–4. Epub 2012/01/06. doi: 10.1056/NEJMc1113592. PubMed PMID: 22216764; PubMed Central PMCID: PMC3513272.

17. Haeusler AR, Donnelly CJ, Periz G, Simko EA, Shaw PG, Kim MS, et al. C9orf72 nucleotide repeat structures initiate molecular cascades of disease. Nature. 2014;507(7491):195–200. Epub 2014/03/07. doi: nature13124 [pii] 10.1038/nature13124. PubMed PMID: 24598541; PubMed Central PMCID: PMC4046618.

18. Maizels N. G4-associated human diseases. EMBO Rep. 2015;16(8):910–22. Epub 2015/07/08. doi: embr.201540607 [pii] 10.15252/embr.201540607. PubMed PMID: 26150098; PubMed Central PMCID: PMC4552485.

19. Lopes J, Piazza A, Bermejo R, Kriegsman B, Colosio A, Teulade-Fichou MP, et al. G-quadruplex-induced instability during leading-strand replication. EMBO J. 2011;30(19):4033–46. Epub 2011/08/30. doi: emboj2011316 [pii] 10.1038/emboj.2011.316. PubMed PMID: 21873979; PubMed Central PMCID: PMC3209785.

20. Piazza A, Serero A, Boule JB, Legoix-Ne P, Lopes J, Nicolas A. Stimulation of gross chromosomal rearrangements by the human CEB1 and CEB25 minisatellites in Saccharomyces cerevisiae depends on G-quadruplexes or Cdc13. PLoS Genet. 2012;8(11):e1003033. Epub 2012/11/08. doi: 10.1371/journal.pgen.1003033 PGENETICS-D-12-01421 [pii]. PubMed PMID: 23133402; PubMed Central PMCID: PMC3486850.

21. Kim N, Jinks-Robertson S. Guanine repeat-containing sequences confer transcription-dependent instability in an orientation-specific manner in yeast. DNA Repair (Amst). 2011;10(9):953–60. Epub 2011/08/05. doi: S1568-7864(11)00171-6 [pii] 10.1016/j.dnarep.2011.07.002. PubMed PMID: 21813340; PubMed Central PMCID: PMC3162091.

22. Yadav P, Harcy V, Argueso JL, Dominska M, Jinks-Robertson S, Kim N. Topoisomerase I plays a critical role in suppressing genome instability at a highly transcribed G-quadruplex-forming sequence. PLoS Genet. 2014;10(12):e1004839. Epub 2014/12/05. doi: 10.1371/journal.pgen.1004839 PGENETICS-D-14-02147 [pii]. PubMed PMID: 25473964; PubMed Central PMCID: PMC4256205.

23. Brazda V, Haronikova L, Liao JC, Fojta M. DNA and RNA quadruplex-binding proteins. Int J Mol Sci. 2014;15(10):17493–517. Epub 2014/10/01. doi: ijms151017493 [pii] 10.3390/ijms151017493. PubMed PMID: 25268620; PubMed Central PMCID: PMC4227175.

24. Pedroso IM, Hayward W, Fletcher TM. The effect of the TRF2 N-terminal and TRFH regions on telomeric G-quadruplex structures. Nucleic Acids Res. 2009;37(5):1541–54. Epub 2009/01/14. doi: gkn1081 [pii] 10.1093/nar/gkn1081. PubMed PMID: 19139067; PubMed Central PMCID: PMC2655686.

25. Bhattacharjee A, Wang Y, Diao J, Price CM. Dynamic DNA binding, junction recognition and G4 melting activity underlie the telomeric and genome-wide roles of human CST. Nucleic Acids Res. 2017;45(21):12311–24. Epub 2017/10/19. doi: 4428939 [pii] 10.1093/nar/gkx878. PubMed PMID: 29040642.

26. Li QJ, Tong XJ, Duan YM, Zhou JQ. Characterization of the intramolecular G-quadruplex promoting activity of Est1. FEBS Lett. 2013;587(6):659–65. Epub 2013/02/05. doi: S0014-5793(13)00055-0 [pii] 10.1016/j.febslet.2013.01.024. PubMed PMID: 23376615.

27. Mendoza O, Bourdoncle A, Boule JB, Brosh RM, Jr., Mergny JL. G-quadruplexes and helicases. Nucleic Acids Res. 2016;44(5):1989–2006. Epub 2016/02/18. doi: gkw079 [pii] 10.1093/nar/gkw079. PubMed PMID: 26883636; PubMed Central PMCID: PMC4797304.

28. Cogoi S, Shchekotikhin AE, Xodo LE. HRAS is silenced by two neighboring G-quadruplexes and activated by MAZ, a zinc-finger transcription factor with DNA unfolding property. Nucleic Acids Res. 2014;42(13):8379–88. Epub 2014/07/12. doi: gku574 [pii] 10.1093/nar/gku574. PubMed PMID: 25013182; PubMed Central PMCID: PMC4117790.

29. Gallo A, Lo Sterzo C, Mori M, Di Matteo A, Bertini I, Banci L, et al. Structure of nucleophosmin DNA-binding domain and analysis of its complex with a G-quadruplex sequence from the c-MYC promoter. J Biol Chem. 2012;287(32):26539–48. Epub 2012/06/19. doi: M112.371013 [pii] 10.1074/jbc.M112.371013. PubMed PMID: 22707729; PubMed Central PMCID: PMC3410995.

30. Cantor SB, Bell DW, Ganesan S, Kass EM, Drapkin R, Grossman S, et al. BACH1, a novel helicase-like protein, interacts directly with BRCA1 and contributes to its DNA repair function. Cell. 2001;105(1):149–60. Epub 2001/04/13. doi: S0092-8674(01)00304-X [pii]. PubMed PMID: 11301010.

31. Seal S, Thompson D, Renwick A, Elliott A, Kelly P, Barfoot R, et al. Truncating mutations in the Fanconi anemia J gene BRIP1 are low-penetrance breast cancer susceptibility alleles. Nat Genet. 2006;38(11):1239–41. Epub 2006/10/13. doi: ng1902 [pii] 10.1038/ng1902. PubMed PMID: 17033622.

32. Wu Y, Shin-ya K, Brosh RM, Jr. FANCJ helicase defective in Fanconia anemia and breast cancer unwinds G-quadruplex DNA to defend genomic stability. Mol Cell Biol. 2008;28(12):4116–28. Epub 2008/04/23. doi: MCB.02210-07 [pii] 10.1128/MCB.02210-07. PubMed PMID: 18426915; PubMed Central PMCID: PMC2423121.

33. Sun H, Karow JK, Hickson ID, Maizels N. The Bloom's syndrome helicase unwinds G4 DNA. J Biol Chem. 1998;273(42):27587–92. Epub 1998/10/09. PubMed PMID: 9765292.

34. Lopez CR, Singh S, Hambarde S, Griffin WC, Gao J, Chib S, et al. Yeast Sub1 and human PC4 are G-quadruplex binding proteins that suppress genome instability at co-transcriptionally formed G4 DNA. Nucleic Acids Res. 2017;45(10):5850–62. Epub 2017/04/04. doi: 3078565 [pii] 10.1093/nar/gkx201. PubMed PMID: 28369605; PubMed Central PMCID: PMC5449603.

35. Tajrishi MM, Tuteja R, Tuteja N. Nucleolin: The most abundant multifunctional phosphoprotein of nucleolus. Commun Integr Biol. 2011;4(3):267–75. Epub 2011/10/08. doi: 10.4161/cib.4.3.148841942-0889-4-3-7 [pii]. PubMed PMID: 21980556; PubMed Central PMCID: PMC3187884.

36. Berger CM, Gaume X, Bouvet P. The roles of nucleolin subcellular localization in cancer. Biochimie. 2015;113:78–85. Epub 2015/04/14. doi: S0300-9084(15)00102-9 [pii] 10.1016/j.biochi.2015.03.023. PubMed PMID: 25866190.

37. Otake Y, Soundararajan S, Sengupta TK, Kio EA, Smith JC, Pineda-Roman M, et al. Overexpression of nucleolin in chronic lymphocytic leukemia cells induces stabilization of bcl2 mRNA. Blood. 2007;109(7):3069–75. Epub 2006/12/21. doi: blood-2006-08-043257 [pii] 10.1182/blood-2006-08-043257. PubMed PMID: 17179226; PubMed Central PMCID: PMC1852223.

38. Satake Y, Kuwano Y, Nishikawa T, Fujita K, Saijo S, Itai M, et al. Nucleolin facilitates nuclear retention of an ultraconserved region containing TRA2beta4 and accelerates colon cancer cell growth. Oncotarget. 2018;9(42):26817–33. Epub 2018/06/22. doi: 10.18632/oncotarget.25510 25510 [pii]. PubMed PMID: 29928487; PubMed Central PMCID: PMC6003563.

39. Gonzalez V, Guo K, Hurley L, Sun D. Identification and characterization of nucleolin as a c-myc G-quadruplex-binding protein. J Biol Chem. 2009;284(35):23622–35. Epub 2009/07/08. doi: M109.018028 [pii] 10.1074/jbc.M109.018028. PubMed PMID: 19581307; PubMed Central PMCID: PMC2749137.

40. Gonzalez V, Hurley LH. The C-terminus of nucleolin promotes the formation of the c-MYC G-quadruplex and inhibits c-MYC promoter activity. Biochemistry. 2010;49(45):9706–14. Epub 2010/10/12. doi: 10.1021/bi100509s. PubMed PMID: 20932061; PubMed Central PMCID: PMC2976822.

41. Tosoni E, Frasson I, Scalabrin M, Perrone R, Butovskaya E, Nadai M, et al. Nucleolin stabilizes G-quadruplex structures folded by the LTR promoter and silences HIV-1 viral transcription. Nucleic Acids Res. 2015;43(18):8884–97. Epub 2015/09/12. doi: gkv897 [pii] 10.1093/nar/gkv897. PubMed PMID: 26354862; PubMed Central PMCID: PMC4605322.

42. Dempsey LA, Sun H, Hanakahi LA, Maizels N. G4 DNA binding by LR1 and its subunits, nucleolin and hnRNP D, A role for G-G pairing in immunoglobulin switch recombination. J Biol Chem. 1999;274(2):1066–71. Epub 1999/01/05. PubMed PMID: 9873052.

43. Erard MS, Belenguer P, Caizergues-Ferrer M, Pantaloni A, Amalric F. A major nucleolar protein, nucleolin, induces chromatin decondensation by binding to histone H1. Eur J Biochem. 1988;175(3):525–30. Epub 1988/08/15. PubMed PMID: 3409881.

44. Ghosh M, Singh M. RGG-box in hnRNPA1 specifically recognizes the telomere G-quadruplex DNA and enhances the G-quadruplex unfolding ability of UP1 domain. Nucleic Acids Res. 2018. Epub 2018/09/25. doi: 10.1093/nar/gky854. PubMed PMID: 30247678.

45. Hanakahi LA, Sun H, Maizels N. High affinity interactions of nucleolin with G-G-paired rDNA. J Biol Chem. 1999;274(22):15908–12. Epub 1999/05/21. PubMed PMID: 10336496.

46. Kondo K, Inouye M. Yeast NSR1 protein that has structural similarity to mammalian nucleolin is involved in pre-rRNA processing. J Biol Chem. 1992;267(23):16252–8. Epub 1992/08/15. PubMed PMID: 1644811.

47. Lee WC, Zabetakis D, Melese T. NSR1 is required for pre-rRNA processing and for the proper maintenance of steady-state levels of ribosomal subunits. Mol Cell Biol. 1992;12(9):3865–71. Epub 1992/09/01. doi: 10.1128/mcb.12.9.3865. PubMed PMID: 1508189; PubMed Central PMCID: PMCPMC360260.

48. Lee WC, Xue ZX, Melese T. The NSR1 gene encodes a protein that specifically binds nuclear localization sequences and has two RNA recognition motifs. J Cell Biol. 1991;113(1):1–12. Epub 1991/04/01. PubMed PMID: 1706724; PubMed Central PMCID: PMC2288927.

49. Yadav P, Owiti N, Kim N. The role of topoisomerase I in suppressing genome instability associated with a highly transcribed guanine-rich sequence is not restricted to preventing RNA:DNA hybrid accumulation. Nucleic Acids Res. 2016;44(2):718–29. doi: 10.1093/nar/gkv1152. PubMed PMID: 26527723; PubMed Central PMCID: PMCPMC4737143.

50. Megonigal MD, Fertala J, Bjornsti MA. Alterations in the catalytic activity of yeast DNA topoisomerase I result in cell cycle arrest and cell death. J Biol Chem. 1997;272(19):12801–8. Epub 1997/05/09. PubMed PMID: 9139740.

51. Wahba L, Amon JD, Koshland D, Vuica-Ross M. RNase H and multiple RNA biogenesis factors cooperate to prevent RNA:DNA hybrids from generating genome instability. Mol Cell. 2011;44(6):978–88. Epub 2011/12/27. doi: S1097-2765(11)00893-8 [pii] 10.1016/j.molcel.2011.10.017. PubMed PMID: 22195970; PubMed Central PMCID: PMC3271842.

52. Chen C, Kolodner RD. Gross chromosomal rearrangements in Saccharomyces cerevisiae replication and recombination defective mutants. Nat Genet. 1999;23(1):81–5. Epub 1999/09/02. doi: 10.1038/12687. PubMed PMID: 10471504.

53. Williams JD, Fleetwood S, Berroyer A, Kim N, Larson ED. Sites of instability in the human TCF3 (E2A) gene adopt G-quadruplex DNA structures in vitro. Front Genet. 2015;6:177. Epub 2015/06/02. doi: 10.3389/fgene.2015.00177. PubMed PMID: 26029241; PubMed Central PMCID: PMC4426816.

54. Hansel-Hertsch R, Beraldi D, Lensing SV, Marsico G, Zyner K, Parry A, et al. G-quadruplex structures mark human regulatory chromatin. Nat Genet. 2016;48(10):1267–72. Epub 2016/09/13. doi: 10.1038/ng.3662. PubMed PMID: 27618450.

55. Soundararajan S, Chen W, Spicer EK, Courtenay-Luck N, Fernandes DJ. The nucleolin targeting aptamer AS1411 destabilizes Bcl-2 messenger RNA in human breast cancer cells. Cancer Res. 2008;68(7):2358–65. doi: 10.1158/0008-5472.CAN-07-5723. PubMed PMID: 18381443.

56. Peled JU, Kuang FL, Iglesias-Ussel MD, Roa S, Kalis SL, Goodman MF, et al. The biochemistry of somatic hypermutation. Annu Rev Immunol. 2008;26:481–511. Epub 2008/02/29. doi: 10.1146/annurev.immunol.26.021607.090236. PubMed PMID: 18304001.

57. Pinheiro LB, Coleman VA, Hindson CM, Herrmann J, Hindson BJ, Bhat S, et al. Evaluation of a droplet digital polymerase chain reaction format for DNA copy number quantification. Anal Chem. 2012;84(2):1003–11. Epub 2011/11/30. doi: 10.1021/ac202578x. PubMed PMID: 22122760; PubMed Central PMCID: PMCPMC3260738.

58. Batrakou DG, Heron ED, Nieduszynski CA. Rapid high-resolution measurement of DNA replication timing by droplet digital PCR. Nucleic Acids Res. 2018;46(19):e112. Epub 2018/07/10. doi: 10.1093/nar/gky590. PubMed PMID: 29986073; PubMed Central PMCID: PMCPMC6212846.

59. Kim N, Abdulovic AL, Gealy R, Lippert MJ, Jinks-Robertson S. Transcription-associated mutagenesis in yeast is directly proportional to the level of gene expression and influenced by the direction of DNA replication. DNA Repair (Amst). 2007;6(9):1285–96. Epub 2007/04/03. doi: 10.1016/j.dnarep.2007.02.023. PubMed PMID: 17398168; PubMed Central PMCID: PMCPMC2034516.

60. Huber MD, Lee DC, Maizels N. G4 DNA unwinding by BLM and Sgs1p: substrate specificity and substrate-specific inhibition. Nucleic Acids Res. 2002;30(18):3954–61. Epub 2002/09/18. PubMed PMID: 12235379; PubMed Central PMCID: PMC137114.

61. Paeschke K, Bochman ML, Garcia PD, Cejka P, Friedman KL, Kowalczykowski SC, et al. Pif1 family helicases suppress genome instability at G-quadruplex motifs. Nature. 2013;497(7450):458–62. Epub 2013/05/10. doi: nature12149 [pii] 10.1038/nature12149. PubMed PMID: 23657261; PubMed Central PMCID: PMC3680789.

62. Indig FE, Rybanska I, Karmakar P, Devulapalli C, Fu H, Carrier F, et al. Nucleolin inhibits G4 oligonucleotide unwinding by Werner helicase. PLoS One. 2012;7(6):e35229. Epub 2012/06/08. doi: 10.1371/journal.pone.0035229 PONE-D-10-04490 [pii]. PubMed PMID: 22675465; PubMed Central PMCID: PMC3366963.

63. Goldstein M, Derheimer FA, Tait-Mulder J, Kastan MB. Nucleolin mediates nucleosome disruption critical for DNA double-strand break repair. Proc Natl Acad Sci U S A. 2013;110(42):16874–9. Epub 2013/10/02. doi: 10.1073/pnas.1306160110. PubMed PMID: 24082117; PubMed Central PMCID: PMCPMC3801049.

64. Pennaneach V, Putnam CD, Kolodner RD. Chromosome healing by de novo telomere addition in Saccharomyces cerevisiae. Mol Microbiol. 2006;59(5):1357–68. Epub 2006/02/14. doi: 10.1111/j.1365-2958.2006.05026.x. PubMed PMID: 16468981.

65. Thandapani P, O’Connor TR, Bailey TL, Richard S. Defining the RGG/RG motif. Mol Cell. 2013;50(5):613–23. doi: 10.1016/j.molcel.2013.05.021. PubMed PMID: 23746349.

66. Ivessa AS, Lenzmeier BA, Bessler JB, Goudsouzian LK, Schnakenberg SL, Zakian VA. The Saccharomyces cerevisiae helicase Rrm3p facilitates replication past nonhistone protein-DNA complexes. Mol Cell. 2003;12(6):1525–36. Epub 2003/12/24. PubMed PMID: 14690605.

67. Azevedo C, Livermore T, Saiardi A. Protein polyphosphorylation of lysine residues by inorganic polyphosphate. Mol Cell. 2015;58(1):71–82. Epub 2015/03/17. doi: S1097-2765(15)00102-1 [pii] 10.1016/j.molcel.2015.02.010. PubMed PMID: 25773596.

68. Han J, van Hoof A. The RNA Exosome Channeling and Direct Access Conformations Have Distinct In Vivo Functions. Cell Rep. 2016;16(12):3348–58. Epub 2016/09/23. doi: 10.1016/j.celrep.2016.08.059. PubMed PMID: 27653695; PubMed Central PMCID: PMCPMC5044803.

69. Lopez CR, Ribes-Zamora A, Indiviglio SM, Williams CL, Haricharan S, Bertuch AA. Ku must load directly onto the chromosome end in order to mediate its telomeric functions. PLoS Genet. 2011;7(8):e1002233. Epub 2011/08/20. doi: 10.1371/journal.pgen.1002233 PGENETICS-D-11-00999 [pii]. PubMed PMID: 21852961; PubMed Central PMCID: PMC3154960.

70. Gao J, Zybailov BL, Byrd AK, Griffin WC, Chib S, Mackintosh SG, et al. Yeast transcription co-activator Sub1 and its human homolog PC4 preferentially bind to G-quadruplex DNA. Chem Commun (Camb). 2015;51(33):7242–4. Epub 2015/03/31. doi: 10.1039/c5cc00742a. PubMed PMID: 25813861; PubMed Central PMCID: PMC4465349.

71. Owiti N, Wei S, Bhagwat AS, Kim N. Unscheduled DNA synthesis leads to elevated uracil residues at highly transcribed genomic loci in Saccharomyces cerevisiae. PLoS Genet. 2018;14(7):e1007516. Epub 2018/07/18. doi: 10.1371/journal.pgen.1007516. PubMed PMID: 30016327; PubMed Central PMCID: PMCPMC6063437.

