## Supplementary figures and images for "High Affinity binding of Yeast Nucleolin Nsr1 to Co-transcriptionally Formed G4 DNA Obstructs Replication and Elevates Genome Instability"

### supplemental figures

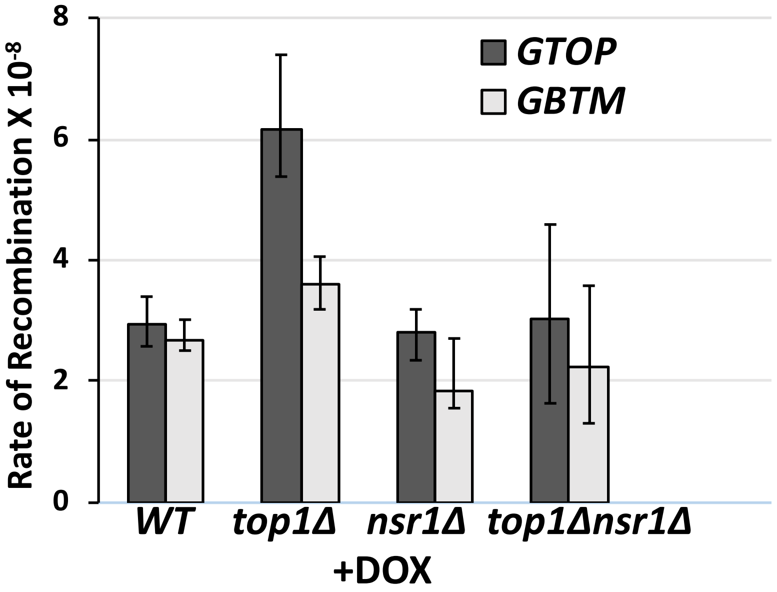

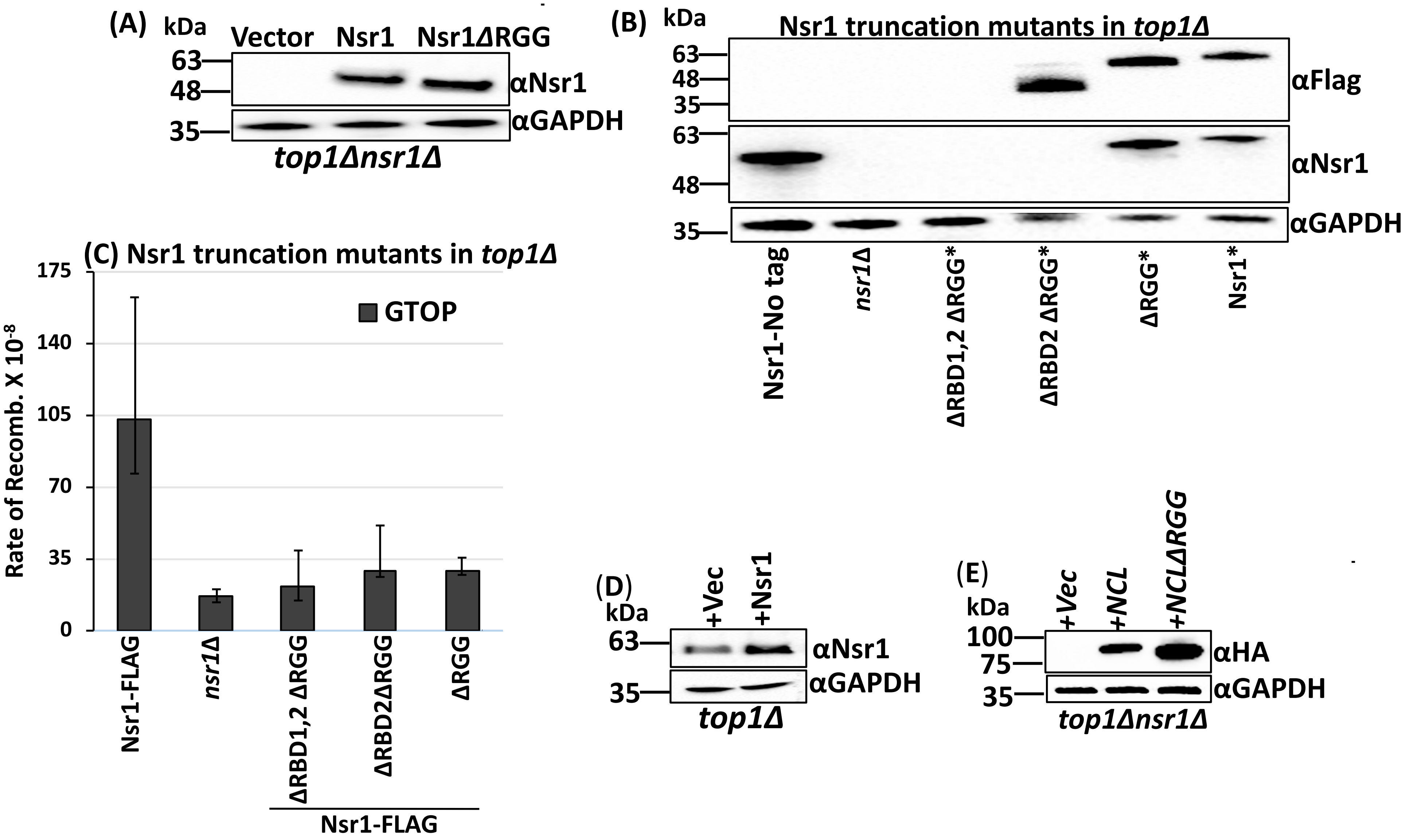

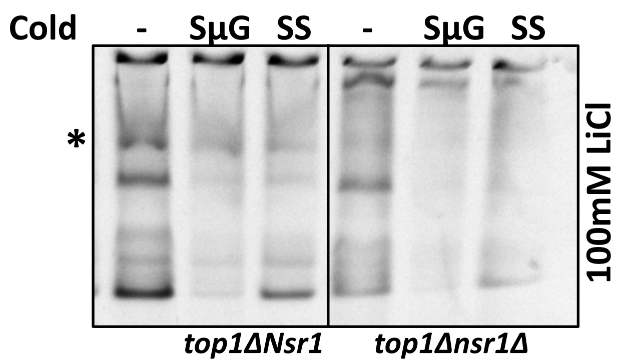

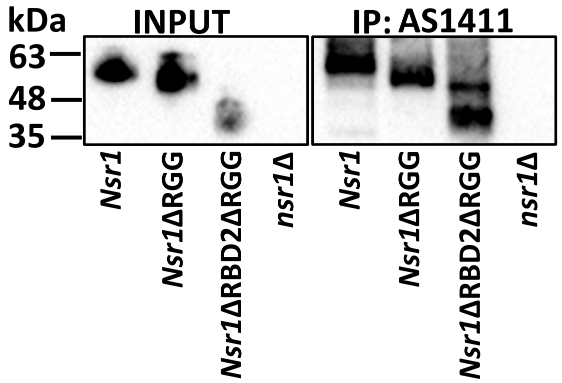

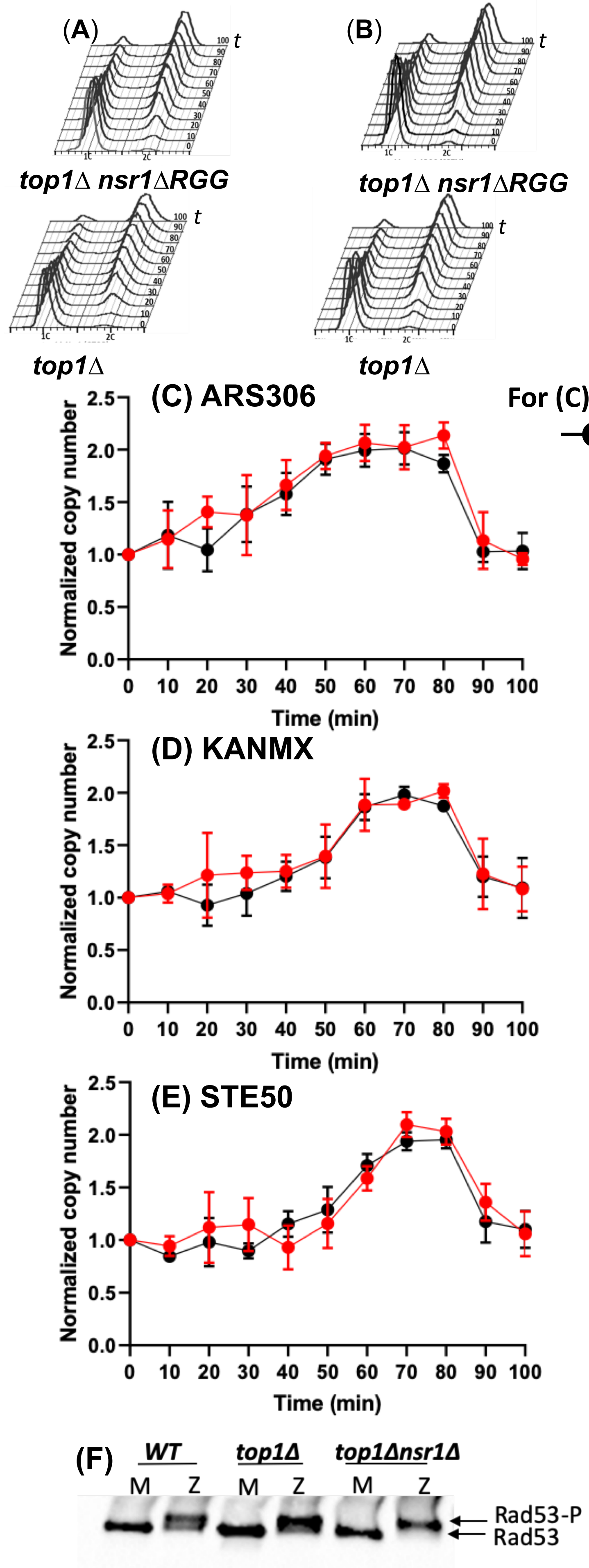
